# Landscape analysis of escape variants identifies SARS-CoV-2 spike mutations that attenuate monoclonal and serum antibody neutralization

**DOI:** 10.1101/2020.11.06.372037

**Authors:** Zhuoming Liu, Laura A. VanBlargan, Louis-Marie Bloyet, Paul W. Rothlauf, Rita E. Chen, Spencer Stumpf, Haiyan Zhao, John M. Errico, Elitza S. Theel, Mariel J. Liebeskind, Brynn Alford, William J. Buchser, Ali H. Ellebedy, Daved H. Fremont, Michael S. Diamond, Sean P. J. Whelan

## Abstract

Although neutralizing antibodies against the SARS-CoV-2 spike (S) protein are a goal of COVID-19 vaccines and have received emergency use authorization as therapeutics, viral escape mutants could compromise their efficacy. To define the immune-selected mutational landscape in S protein, we used a VSV-eGFP-SARS-CoV-2-S chimeric virus and 19 neutralizing monoclonal antibodies (mAbs) against the receptor-binding domain (RBD) to generate 50 different escape mutants. The variants were mapped onto the RBD structure and evaluated for cross-resistance to mAbs and convalescent human sera. Each mAb had a unique resistance profile, although many shared residues within an epitope. Some variants (*e.g*., S477N) were resistant to neutralization by multiple mAbs, whereas others (*e.g*., E484K) escaped neutralization by convalescent sera, suggesting some humans induce a narrow repertoire of neutralizing antibodies. Comparing the antibody-mediated mutational landscape in S with sequence variation in circulating SARS-CoV-2, we define substitutions that may attenuate neutralizing immune responses in some humans.

## INTRODUCTION

Control of the ongoing SARS-CoV-2 pandemic will require deployment of multiple countermeasures including therapeutics and vaccines. Therapeutic candidates that have received emergency use authorization (EUA) or are in development include several monoclonal antibodies (mAbs) (Chen et al., 2020; Group et al., 2020; Weinreich et al., 2020) that recognize the SARS-CoV-2 spike (S) protein, which decorates the virion surface (Ke et al., 2020). The S protein is comprised of an N-terminal subunit (S1) that mediates receptor binding and a C-terminal subunit (S2) responsible for virus-cell membrane fusion (Wrapp et al., 2020). During viral entry into cells, the receptor-binding domain (RBD) of S1 engages the primary receptor, human angiotensin converting enzyme 2 (hACE2) (Letko et al., 2020). Processing of S by host cell proteases, typically TMPRSS2, TMPRSS4, or endosomal cathepsins, facilitates the S2-dependent fusion of viral and host-cell membranes (Hoffmann et al., 2020; Zang et al., 2020). Potently neutralizing antibodies against SARS-CoV-2 target the RBD (Brouwer et al., 2020; Group et al., 2020; Rogers et al., 2020; Wu et al., 2020b; Zost et al., 2020) with many inhibiting infection by blocking receptor engagement (Alsoussi et al., 2020; Wu et al., 2020b). Understanding the epitopes recognized by protective antibodies and whether natural variation in the S protein is associated with resistance to neutralization may predict the utility of antibody-based countermeasures.

RNA viruses exist as a swarm or “quasispecies” of genome sequences around a core consensus sequence (Dolan et al., 2018). Under conditions of selection, such as those imposed by neutralizing antibodies or drugs, variants of the swarm can escape genetically and become resistant. The relative fitness of escape mutants determines whether they are lost rapidly from the swarm or provide a competitive advantage. The intrinsically high error rates of viral RNA-dependent RNA polymerases (RdRp) result in the stochastic introduction of mutations during viral genome replication with substitutions approaching a nucleotide change per genome for each round of replication (Sanjuan et al., 2010). Coronaviruses, because of their large genome size, encode a proofreading 3’-to-5’ exoribonuclease (ExoN, nsp14) that helps to correct errors made by the RdRp during replication (Smith and Denison, 2013). As a result of ExoN activity, the frequency of escape from antibody neutralization by coronaviruses is less than for other RNA viruses lacking such an enzyme (Smith et al., 2013).

To date, 4150 mutations have been identified in the S gene of SARS-CoV-2 isolated from humans (CoV-GLUE, 2021; GISAID, 2021). These mutations give rise to 1,246 amino acid changes including 187 substitutions in the RBD. The abundance of many variants in the human population suggests they are not accompanied by a fitness loss. Multiple mechanisms likely account for the emergence of such substitutions including host adaptation, immune selection during natural infection, and possibly reinfection of individuals with incomplete or waning immunity. Convalescent plasma therapy, vaccination, and administration of therapeutic antibodies each could select for additional variants, and their effectiveness as countermeasures might be compromised by preexisting resistant mutants. Thus, as therapeutic antibodies and vaccines are deployed, it is increasingly important to define the patterns of antibody resistance that arise. The impact of SARS-CoV-2 adaptation for infection of other hosts including mice (Dinnon et al., 2020; Gu et al., 2020), mink (Oude Munnink et al., 2020) and domesticated animals (Halfmann et al., 2020; Shi et al., 2020) could also contribute to selection of new variants.

Here, we used a panel of antibodies including the previously reported 2B04, 1B07 and 2H04 mAbs (Alsoussi et al., 2020) and newly-generated neutralizing mAbs against SARS-CoV-2 RBD to select for escape variants and define the mutational landscape of resistance. To facilitate selection, we used a chimeric, infectious vesicular stomatitis virus (VSV) in which the endogenous glycoprotein was replaced with the S protein of SARS-CoV-2 (Case et al., 2020). VSV-eGFP-SARS-CoV-2-S_∆21_ (herein, VSV-SARS-CoV-2) replicates to high titer (10^7^-10^8^ plaque forming units ml^−1^ within 48 hours), mimics the SARS-CoV-2 requirement for human ACE2 as a receptor, and is neutralized by SARS-CoV-2 S-specific mAbs (Case et al., 2020). In three selection campaigns using 19 different mAbs, we isolated 50 different escape mutants within the RBD. Many escape mutations arose proximal to or within the ACE2 binding footprint suggesting that multiple neutralizing mAbs inhibit infection by interfering with receptor engagement. Cross-neutralization studies involving 29 of the escape mutants and 10 mAbs identified mutants that were resistant to multiple antibodies and also those with unique resistance profiles. Remarkably, substitutions at residue E484 of S protein were associated with resistance to neutralization by polyclonal human immune sera, suggesting that some individuals generate neutralizing antibodies recognizing a focused target on the RBD. Resistance to inhibition by soluble recombinant human ACE2, a candidate decoy molecule drug (Chan et al., 2020; Monteil et al., 2020) currently in clinical trials (NCT04375046, NCT04287686), was observed with an F486S substitution. Cross-referencing of our 50 resistant mutants with sequences of clinical isolates of SARS-CoV-2 demonstrates that some already circulating variants will be resistant to monoclonal and polyclonal antibodies. This data and functional approach may be useful for monitoring and evaluating the emergence of escape from antibody-based therapeutic and vaccine countermeasures as they are deployed.

## RESULTS

### Selection of mAb escape mutants in SARS-CoV-2 S

To select for SARS-CoV-2 S variants that escape neutralization, we used VSV-SARS-CoV-2 (Case et al., 2020) and mAb 2B04, which was generated from cloned murine B cells following immunization of C57BL/6 mice with recombinant RBD and boosted with recombinant S. Antibody neutralization resistant mutants were recovered by plaque isolation (**Fig 1A**), and their resistance was verified by subsequent virus infection in the presence or absence of antibody. Antibody 2B04 failed to inhibit VSV-SARS-CoV-2 resistant variants as judged by plaque number and size (**Fig 1B**). Sequence analysis identified the mutations E484A, E484K, and F486S (**Fig 1B**), each of which fall within the RBD and map to residues involved in ACE2 binding (Lan et al., 2020) (**Fig 2**).

**Figure 1.**
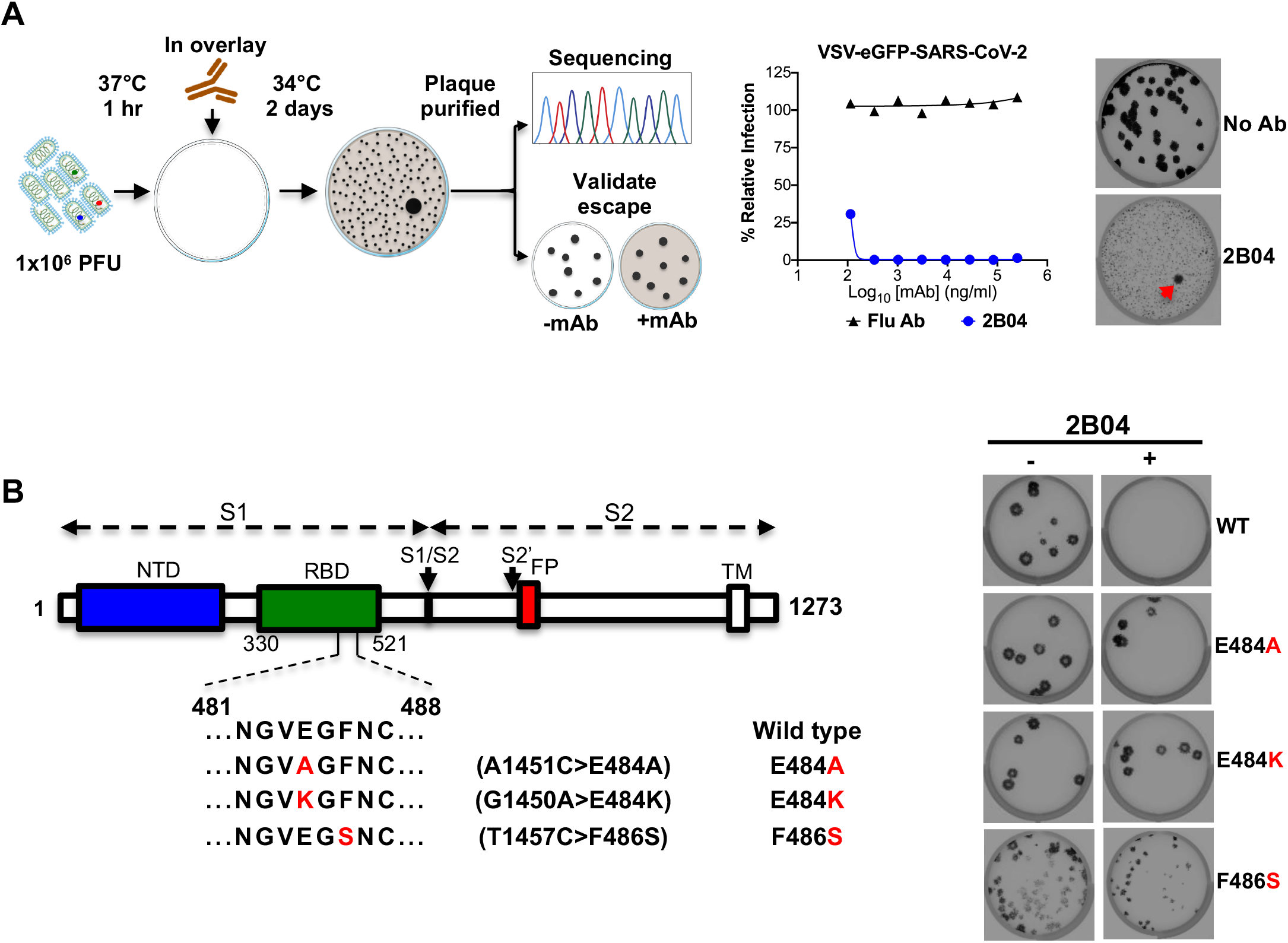
VSV-SARS-CoV-2 escape mutant isolation. (**A**) Outline of escape mutant selection experiment. 2B04 and a control anti-influenza virus mAb were tested for neutralizing activity against VSV-SARS-CoV-2. The concentration of 2B04 added in the overlay completely inhibited viral infection (middle panel). Data are representative of two independent experiments. Plaque assays were performed to isolate the VSV-SARS-CoV-2 escape mutant on Vero E6 TMPRSS2 cells (red arrow indicated). Plaque assays with 2B04 in the overlay (*Bottom plaque in the right panel*); plaque assays without Ab in the overlay (*Top plaque in the right panel*). Data are representative of three independent experiments. (**B**) Schematic of S gene, which underwent Sanger sequencing to identify mutations (*left panel*). For validation, each VSV-SARS-CoV-2 mutant was tested in plaque assays with or without 2B04 in the overlay on Vero cells (*right panel*). Representative images of two independent experiments are shown.

**Figure 2.**
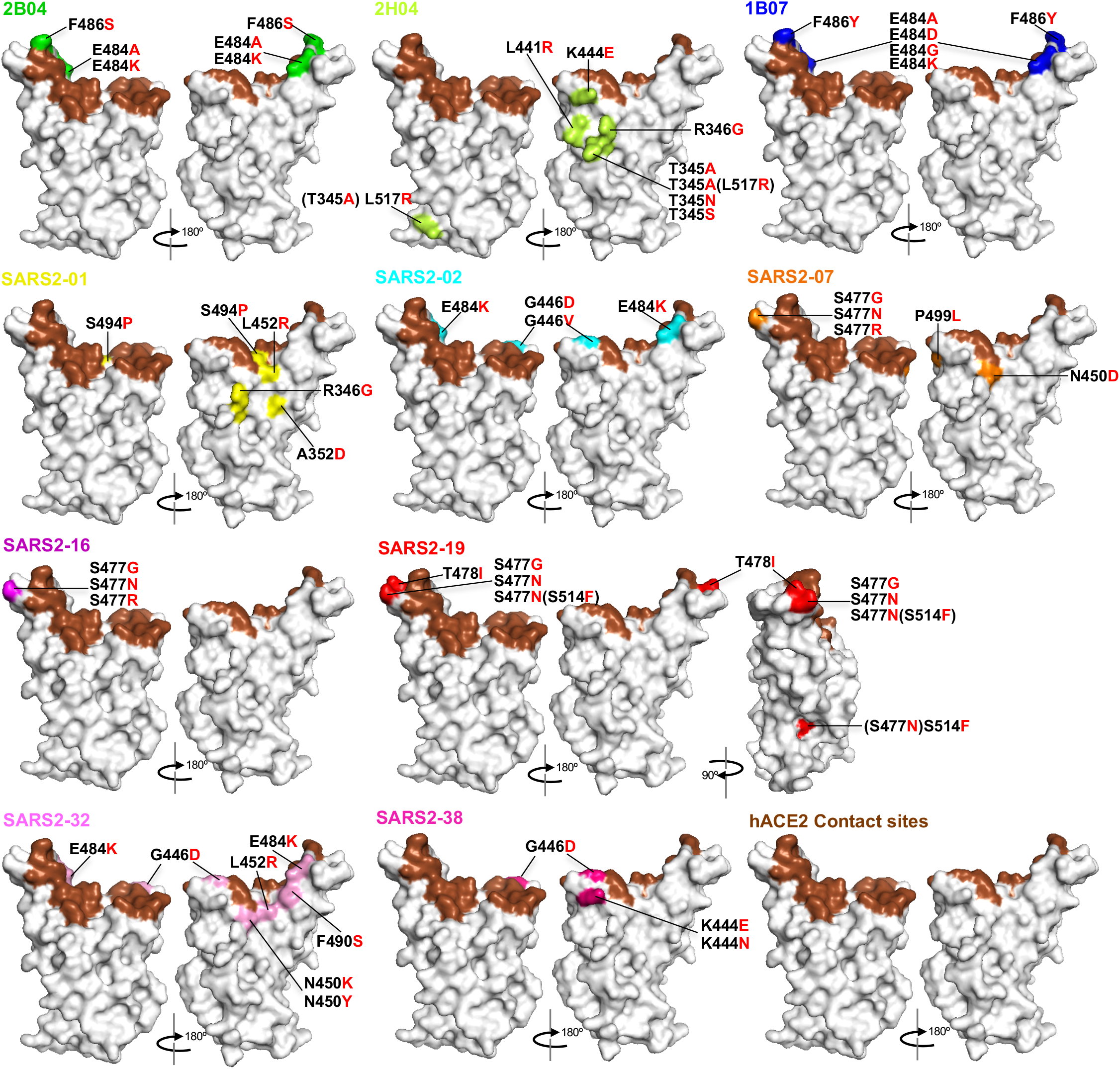
Mapping of escape mutations. The surface model of RBD (from PDB 6M0J) is depicted, and contact residues of the SARS-CoV-2 RBD-hACE2 interfaces are colored in brown. Amino acids whose substitution confers resistance to each mAb in plaque assays are indicated for 2B04 (green), 2H04 (lemon), 1B07 (blue), SARS2-01 (yellow), SARS2-02 (teal), SARS2-07 (tangerine), SARS2-16 (violet), SARS2-19 (red), SARS2-32 (fuschia), and SARS2-38 (magenta). See **Figure S1 and S2**.

We extended this neutralization escape approach to nine additional inhibitory mAbs (**Fig S1, S2 and Table 1**). Sequence analysis of each isolated plaque identified multiple mutations within the RBD (**Table 2**), which we positioned on the reported crystal structure (PDB: 6M0J) (**Fig 2**): 2B04 (green), 2H04 (lime), 1B07 (blue), SARS2-01 (yellow), SARS2-02 (teal), SARS2-07 (tangerine), SARS2-16 (violet), SARS2-19 (red), SARS2-32 (fuschia), and SARS2-38 (magenta). Substitutions that led to resistance of mAbs 2B04, 1B07, SARS2-02, SARS2-07, SARS2-16, SARS2-32, and SARS2-38 cluster within and proximal to the ACE2 binding site. Resistance to antibodies SARS2-01 and SARS2-19 mapped to substitutions at sites on the side of the RBD (**Fig 2**). MAb 2H04 gave rise to resistance mutations that map exclusively on the side and base of the RBD (**Fig 2**). The identification of resistance mutations at the side of the RBD suggest that the mechanism of virus neutralization may be allosteric or possibly through blocking interactions with alternative attachment factors. The presence of resistance mutations at the base of RBD, which lie outside the 2H04 binding footprints, suggests an allosteric mechanism of resistance, perhaps related to the ability of the RBD to adopt the ‘up’ conformation requisite for ACE2 binding.

**Table 1.** Neutralizing mAbs.

| mAb | Immunogen<sup>a</sup> | Isotype | Concentration<br>μg/mL | EC50 ng/mL<sup>b</sup> |
| --- | --- | --- | --- | --- |
| 2B04 | RBD, S, S | Human IgG1<sup>c</sup> | 7,630 | 1.46 |
| 2H04 | RBD, S, S | Human IgG1<sup>c</sup> | 6,700 | 279.3 |
| 1B07 | RBD, S, S | Human IgG1<sup>c</sup> | 5,200 | 15 |
| SARS2-01 | RBD, S, RBD | mouse IgG1 | 44 | 86 |
| SARS2-02 | RBD, S, RBD | mouse IgG1 | 40 | 7 |
| SARS2-07 | RBD, S, RBD | mouse IgG1 | 87 | 65 |
| SARS2-16 | RBD, S, S | mouse IgG1 | 111 | 27 |
| SARS2-19 | RBD, S, S | mouse IgG1 | 31 | 15 |
| SARS2-32 | RBD, S, RBD | mouse IgG1 | 50 | 23 |
| SARS2-38 | RBD, S, S | mouse IgG1 | 49 | 8 |
| SARS2-21 | RBD, S, RBD | mouse IgA | 42 | 24 |
| SARS2-22 | RBD, S, S | mouse IgG1 | 25 | 10 |
| SARS2-23 | RBD, S, S | mouse IgG1 | 126 | 65 |
| SARS2-31 | RBD, S, S | mouse IgG1 | 36 | 28 |
| SARS2-34 | RBD, S, S | mouse IgG1 | 40 | 19 |
| SARS2-55 | RBD, S, RBD | mouse IgG1 | 37 | 10 |
| SARS2-58 | RBD, S, S | mouse IgG1 | 27 | 6 |
| SARS2-66 | RBD, S, S | mouse IgG1 | 44 | 12 |
| SARS2-71 | RBD, S, S | mouse IgG1 | 49 | 11 |

<sup>a</sup> The order of immunogens used to immunize the mice, as described in the Methods.
<sup>b</sup> Neutralization of SARS-CoV-2 by each mAb was assessed by focus-reduction neutralization test. The half-maximal effective concentration (EC50 value) was determined by nonlinear regression. Results are the geometric mean from three to four independent experiments.
<sup>c</sup> mAb was identified as mouse IgG1 and expressed as human IgG1.

**Table 2.**
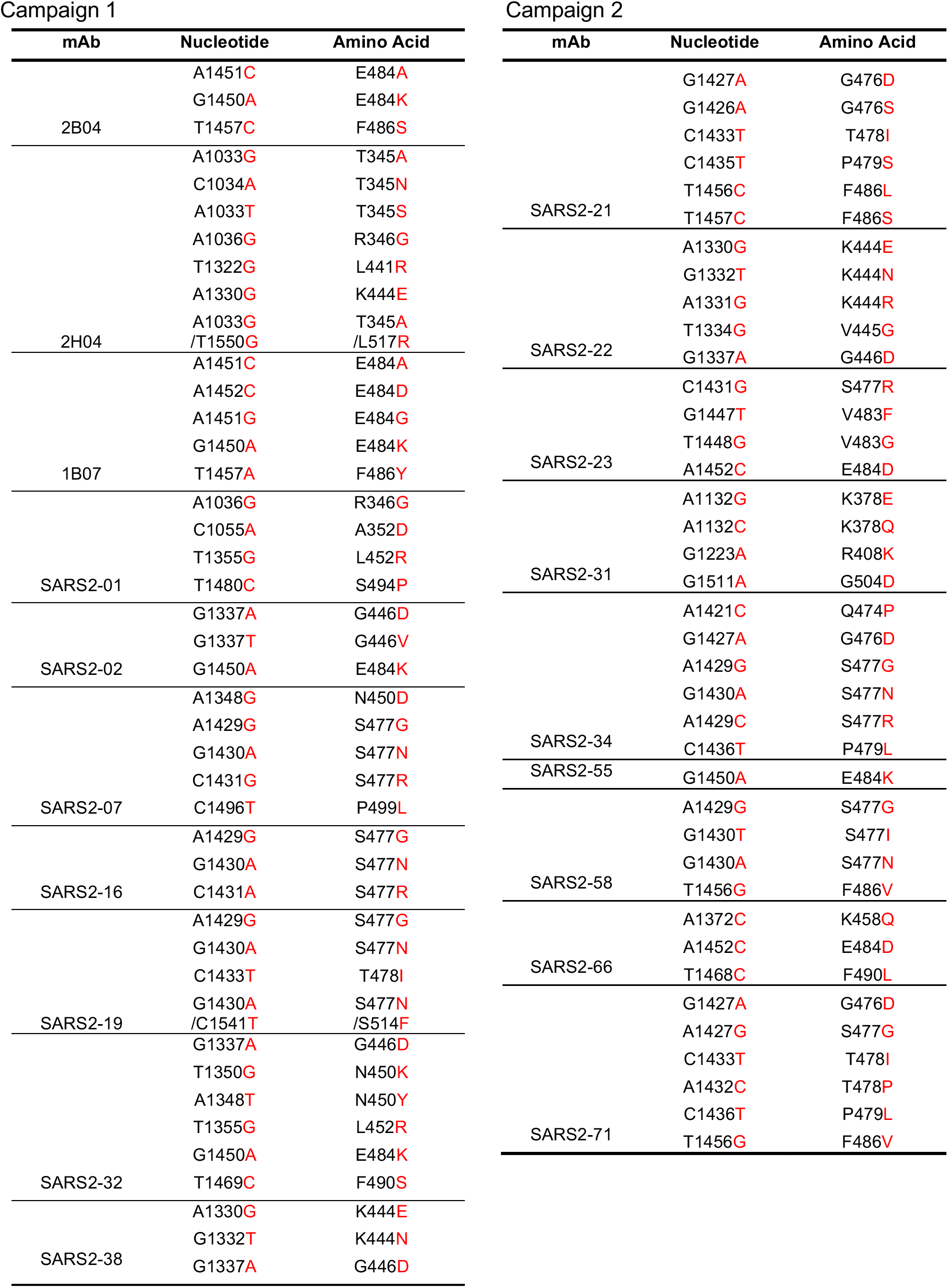
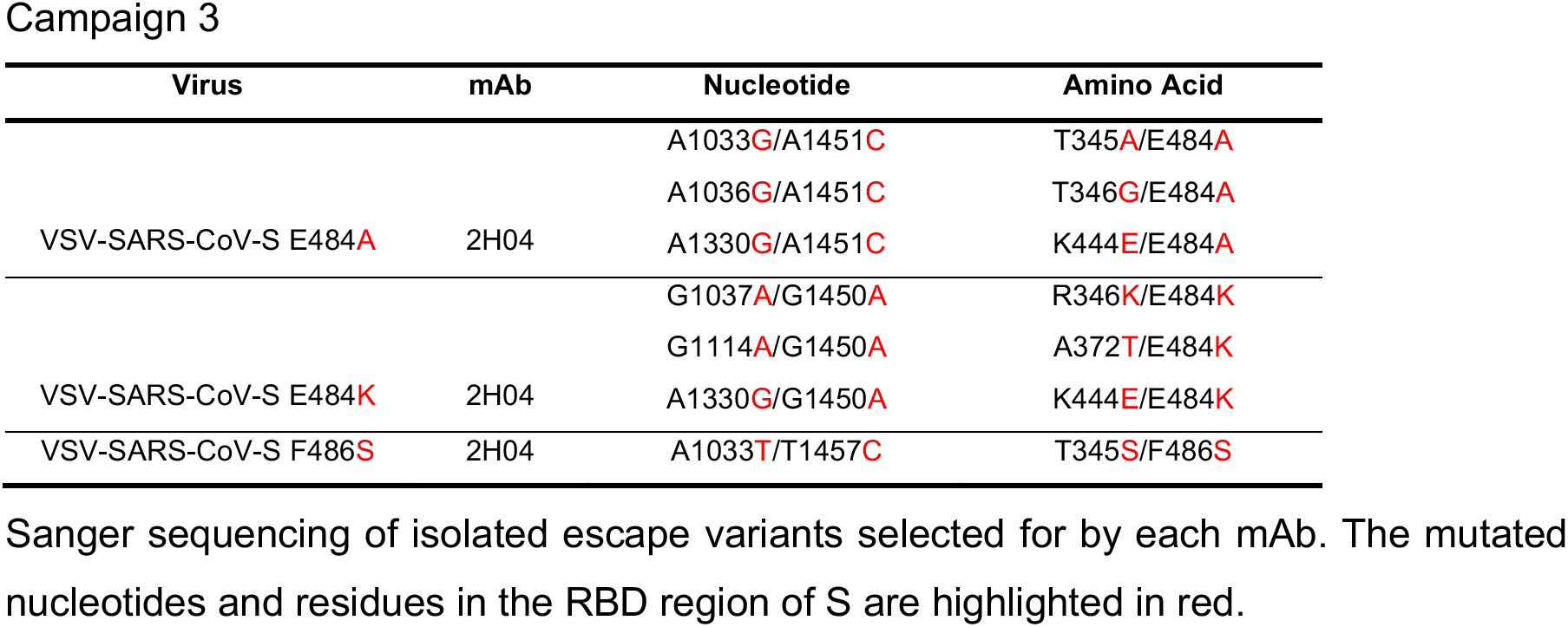
List of mutants.

From this panel of mAbs, we observed resistance substitutions at shared positions. Four mAbs yielded substitutions at E484 (2B04, 1B07, SARS2-02, and SARS2-32), three resulted in changes to residues G446 (SARS2-02, SARS2-32, and SARS2-38) and S477 (SARS2-07, SARS2-16 and SARS2-19), and two promoted escape substitutions at F486 (2B04 and 1B07), K444 (2H04 and SARS2-38), L452 (SARS2-01 and SARS2-32), and N450 (SARS2-07 and SARS2-32), and R346 (2H04 and SARS2-01). The overlapping nature of these epitopes suggests they represent major antigenic sites within the RBD. Although amino acid changes were selected at the same position, many of the substitutions were distinct, consistent with a unique mode of binding for each antibody.

Two mAbs gave rise to variants containing linked amino acid substitutions: 2H04 (T345A and L517R) and SARS2-19 (S477N and S514F). For 2H04, substitution T345A likely arose first, as we isolated this mutation alone, and acquisition of the L517R substitution appeared to enhance infectivity as judged by plaque morphology (**Fig S2**). For SARS2-19, S477N was isolated as a single variant suggesting that this substitution arose first, however acquisition of the S514F did not alter plaque morphology (**Fig S2**). As the L517R or S514F substitutions were not identified in isolation, it remains unclear whether they cause resistance to 2H04 or SARS2-19 respectively. Collectively, these results show that escape mutational profiling can identify key epitopes and dominant antigenic sites.

### Escape mutants confer cross-resistance to multiple mAbs

We next evaluated whether individual mutants could escape neutralization by the other inhibitory mAbs in the panel. We tested the 29 identified escape mutants for neutralization by ten different mAbs. We defined the degree of resistance as a percentage by expressing the number of plaques formed by each mutant in the presence or absence of antibody. We plotted the degree of resistance to neutralization as a heat map and arbitrarily set 50% as the cut-off value for defining resistance (**Fig 3A**). Substitutions at residues T345, R346, K444, G446, N450, L452, S477, T478, E484, F486 and P499 each were associated with resistance to more than one mAb, with substitutions at S477 and E484 exhibiting broad resistance (**Fig 3A**). For residues at which multiple alternate amino acids with different side chains were selected, each particular substitution was associated with a unique resistance profile. For example, K444E was resistant to SARS2-38 and 2H04 with some resistance to SARS2-1, SARS2-2 and SARS2-7, whereas K444N conferred complete resistance to SARS2-38, partial resistance to 2H04 and only weak resistance to SARS2-1 and SARS2-2. G446D was resistant to SARS2-2, SARS2-32 and SARS2-38, but G446V acquired resistance to SARS2-01. Substitutions N450K and N450Y were resistant to SARS2-01 and SARS2-32, whereas N450D facilitated resistance to SARS2-07. Substitution L452R conferred resistance to SARS2-01, SARS2-02, and SARS2-32; S477N, S477G and S477R were each highly resistant to SARS2-07, SARS2-16, and SARS2-19, and S477N and S477G result in a degree of resistance across the entire panel of antibodies; and T478I yielded resistance to SARS2-16 and SARS2-19.

**Figure 3.**
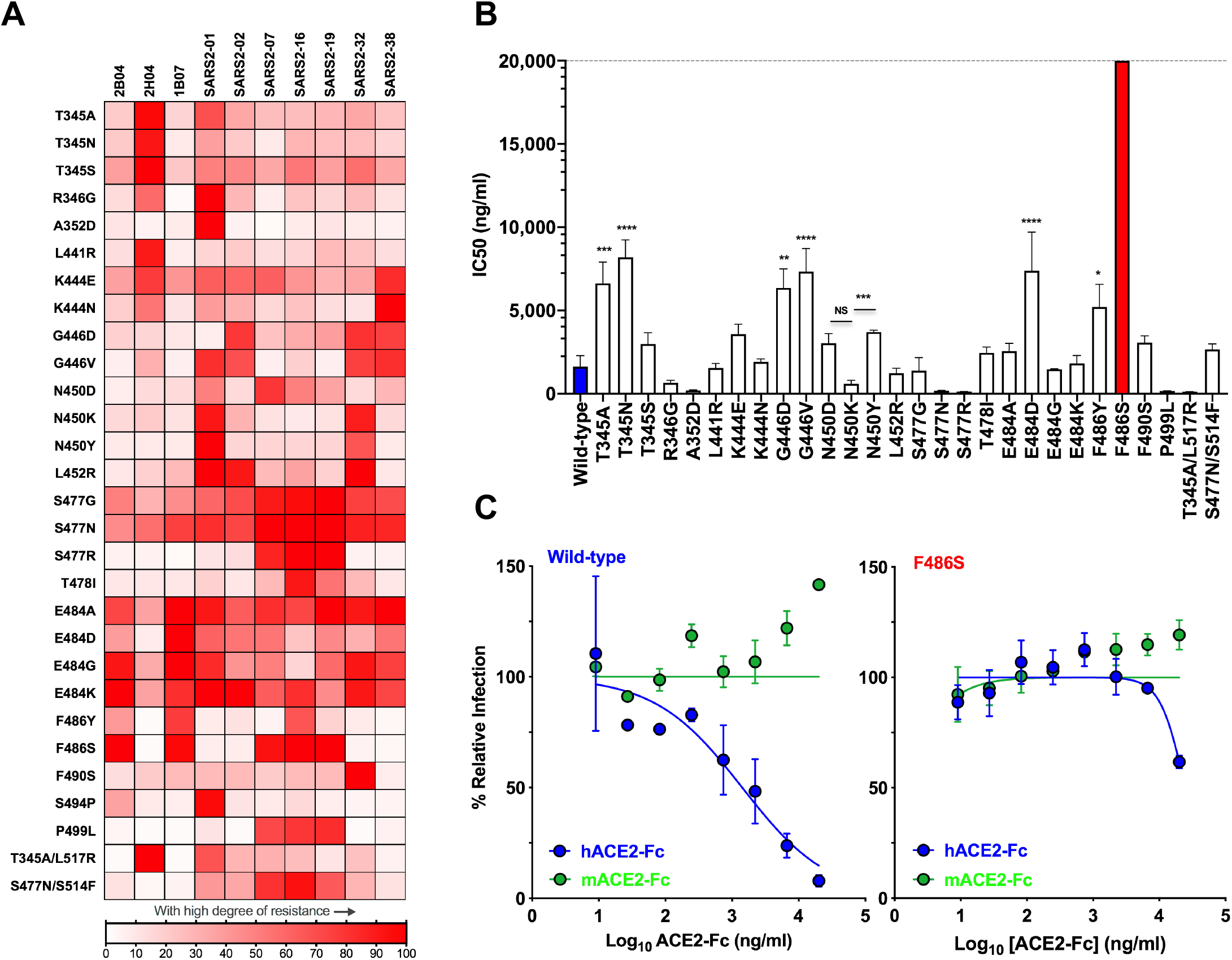
Map of cross-neutralizing activity of VSV-SARS-CoV-2 mutants and neutralization potency of hACE2 decoy receptors against each VSV-SARS-CoV-2 mutant. **(A)** Neutralization of VSV-SARS-CoV-2 mutants was evaluated by plaque assays. Degree of resistance was defined as percentage by expressing the number of plaques formed by each mutant in the presence or absence of antibody versus and is represented as a heatmap from white (low degree of resistance) to red (high degree of resistance). Representative images of two independent experiments are shown in **Figure S3**. (**B**) Neutralization assay of VSV-SARS-CoV-2 mutants in the presence of hACE2-Fc. Virus was incubated with mACE2 or hACE2 at concentrations ranging from 9 ng/ml to 20 μg/ml for 1 h a 37°C and cells were scored for infection at 7.5 h post inoculation by automated microscopy. IC_50_ values were calculated for each virus-hACE2 combination from three independent experiments. (* *P* < 0.05, ** *P* <0.01, *** *P* < 0.001, **** *P* < 0.0001; one-way ANOVA with Dunnett’s post-test; error bars indicate standard error of the mean [SEM]). (**C**) Representative neutralization curves of wild-type and F486S mutant VSV-SARS-CoV-2 with hACE2-Fc and mACE2-Fc. Error bars represent the SEM. Data are representative of three independent experiments. Neutralization curves are provided in **Figure S4**.

Escape variants at residue E484 were isolated using 2B04, 1B07, SARS2-02, and SARS2-32, and specific substitutions at this residue led to varying degrees of resistance across the entire panel of antibodies. E484A exhibited a high degree of resistance to 2B04, 1B07, SARS2-01, SARS2-07, SARS2-19, SARS2-32, SARS2-38; E484G exhibited resistance to 2B04, 1B07, SARS2-01, SARS2-32 and SARS2-38; E484K was resistant to 2B04, 1B07, SARS2-01, SARS2-02, SARS2-16 and SARS2-32; and E484D was resistant only to 1B07 (**Fig 3A and S3**). Substitution F486S was resistant to 2B04, 1B07, SARS2-07, SARS2-16 and SARS2-19, whereas F486Y exhibited resistance only to 1B07 and SARS2-16. Finally, substitution P499L was resistant to SARS2-07, SARS2-16, and SARS2-19. In addition to demonstrating that some mutations confer resistance to multiple neutralizing mAbs, these data suggest that each mAb recognizes distinct yet partially overlapping epitopes.

### Soluble human ACE2-Fc receptor decoy inhibition of escape mutants

Soluble human ACE2 decoy receptors are under evaluation in clinical trials for treatment of COVID-19 (NCT04375046 and NCT04287686). As several of the escape mutants contain substitutions within or proximal to the ACE2 binding site, we evaluated the ability of soluble recombinant ACE2 to inhibit infection of each variant. We incubated each VSV-SARS-CoV-2 mutant with increasing concentrations of soluble human (h) or murine (m) ACE2-Fc for 1 h at 37°C and measured residual infectivity on Vero E6 cells (**Fig 3B and S4**). As observed with chimeric viruses expressing the wild-type S protein, the escape mutants were inhibited by hACE2-Fc but not mACE2-Fc. However, the extent of neutralization by hACE2-Fc varied substantially (**Fig 3B**), with some mutants more sensitive to receptor inhibition and others exhibiting relative resistance. Substitutions at residues R346, A352, N450, S477, S494 and P499 were more sensitive to inhibition by soluble hACE2 than the wild-type S as evidenced by reduced IC_50_ values (**Fig 3B**) and leftward shifts of the inhibition curves (**Fig S4**). This effect was substitution-dependent, as N450K was 6-fold more sensitive to hACE2 than N450Y (*P* < 0.001). Several mutants required higher (3-5-fold) concentrations of hACE2 to block infection, including substitutions at T345A, T345N, G446D, G446V, E484D and F486Y. Again, the specific substitution of a given residue impacted the effect, as T345A and T345N required higher concentrations of hACE2 to inhibit infection, whereas T345S was similar to wild type. Of the 4 substitutions observed at position E484, only E484D was less sensitive (4.6-fold, *P* < 0.0001) to hACE2 inhibition. The most striking effect was observed for F486S, where we achieved only 38% inhibition at the highest concentration (20 μg/ml) of hACE2-Fc tested (**Fig 3B and C**). Residue F486 is located on the top of the hACE2 contact loop of RBD, and the presence of a large hydrophobic residue facilitates efficient receptor engagement (Lan et al., 2020; Shang et al., 2020). Although this substitution alters sensitivity to soluble ACE2 inhibition of infection, its impact on cell surface ACE2 engagement by virus was not examined.

### Escape mutants exhibit resistance to neutralization by polyclonal human immune sera

We previously evaluated the ability of convalescent sera from SARS-CoV-2-infected humans to neutralize VSV-SARS-CoV-2 and defined a strong correlation with inhibition of a clinical isolate of SARS-CoV-2 using a focus reduction neutralization test (FRNT) (Case et al., 2020). We tested four of the serum samples (13, 29, 35 and 37) from patients who had recovered from COVID-19 against our panel of escape mutants (**Table 3**). All four sera neutralized infection of VSV-SARS-CoV-2-S displaying the wild-type S protein as we previously demonstrated. Remarkably, several of the escape mutants were resistant to neutralization at the highest concentration (1:80 initial dilution) of sera tested. All four of the substitutions at residue E484 were resistant to each of the four sera, suggesting that this is part of a dominant neutralizing epitope. Indeed, change at E484 was the only position that led to escape from sera 29 (**Fig 4A-B, and S5A**). Four other substitutions (K444E, G446V, L452R and F490S) resulted in resistance to neutralization of sera 13, 35 and 37 (**Fig 4A and S5A**). Substitutions N450D and N450Y but not N450K were resistant to sera 13 and 35. Sera 13 and 35 also did not efficiently neutralize S477G, L441R, and T478I. All four sera neutralized the single substitution S477N as well as wild-type virus (**Fig 4A-B**). Substitution S477N was sensitive to neutralization by sera 13 and 35 except in the presence of a second S514F substitution (**Fig 4A and S5A**). Additional amino acid substitutions that conferred resistance to serum 13 include T345S and G446D. Substitution F486S, which altered sensitivity to soluble ACE2, escaped neutralization by serum 35 but not 13, 29 or 37. We consistently noticed that some sera also led to an increase in infectivity of specific escape mutants (*e.g*., E484A) at some concentrations (**Fig 4B**). The significance of this increase is unclear, but was observed consistently across sera for several mutants (**Fig S5A**).

**Figure 4.**
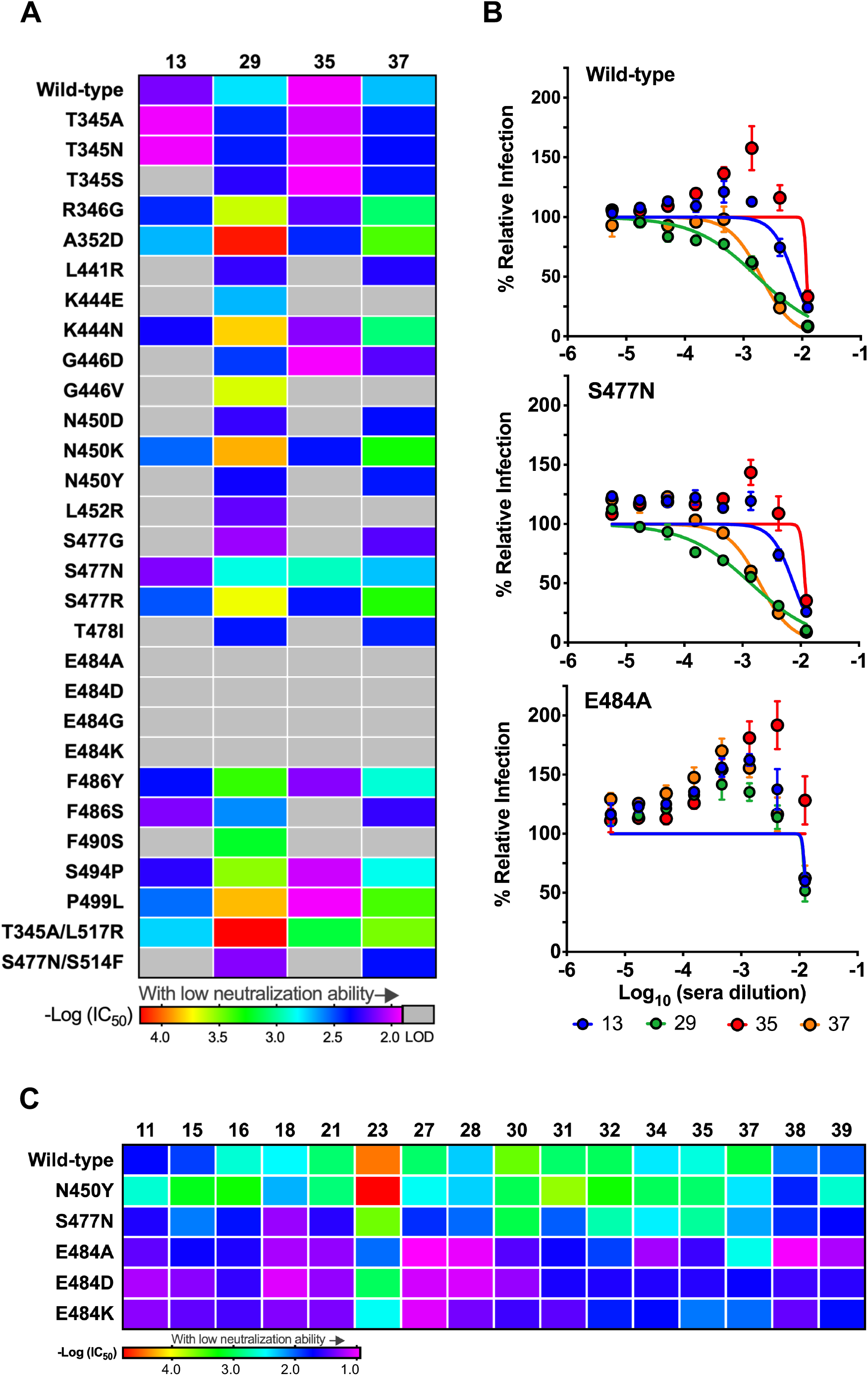
Neutralization potency of human serum against each VSV-SARS-CoV-2 mutant. (**A**) Neutralization potency of four human sera against VSV-SARS-CoV-2 mutants. IC_50_ values were calculated from three independent experiments. Neutralization potency is represented as a rainbow color map from red (most potent with low IC_50_) to violet (less potent with high IC_50_). LOD indicates limit of detection (1:80) (**B**) Representative neutralization curves of wild-type, S477N and E484A mutant with four different human sera. Error bars represent the SEM. Data are representative of three independent experiments. (**C**) Neutralization potency of additional 16 human sera against VSV-SARS-CoV-2 mutants. IC_50_ values were calculated from one independent experiment each. Neutralization potency is represented as a rainbow color map from red (most potent with low IC_50_) to violet (less potent with high IC_50_). Neutralization curves are provided in **Figure S5**.

**Table 3.**
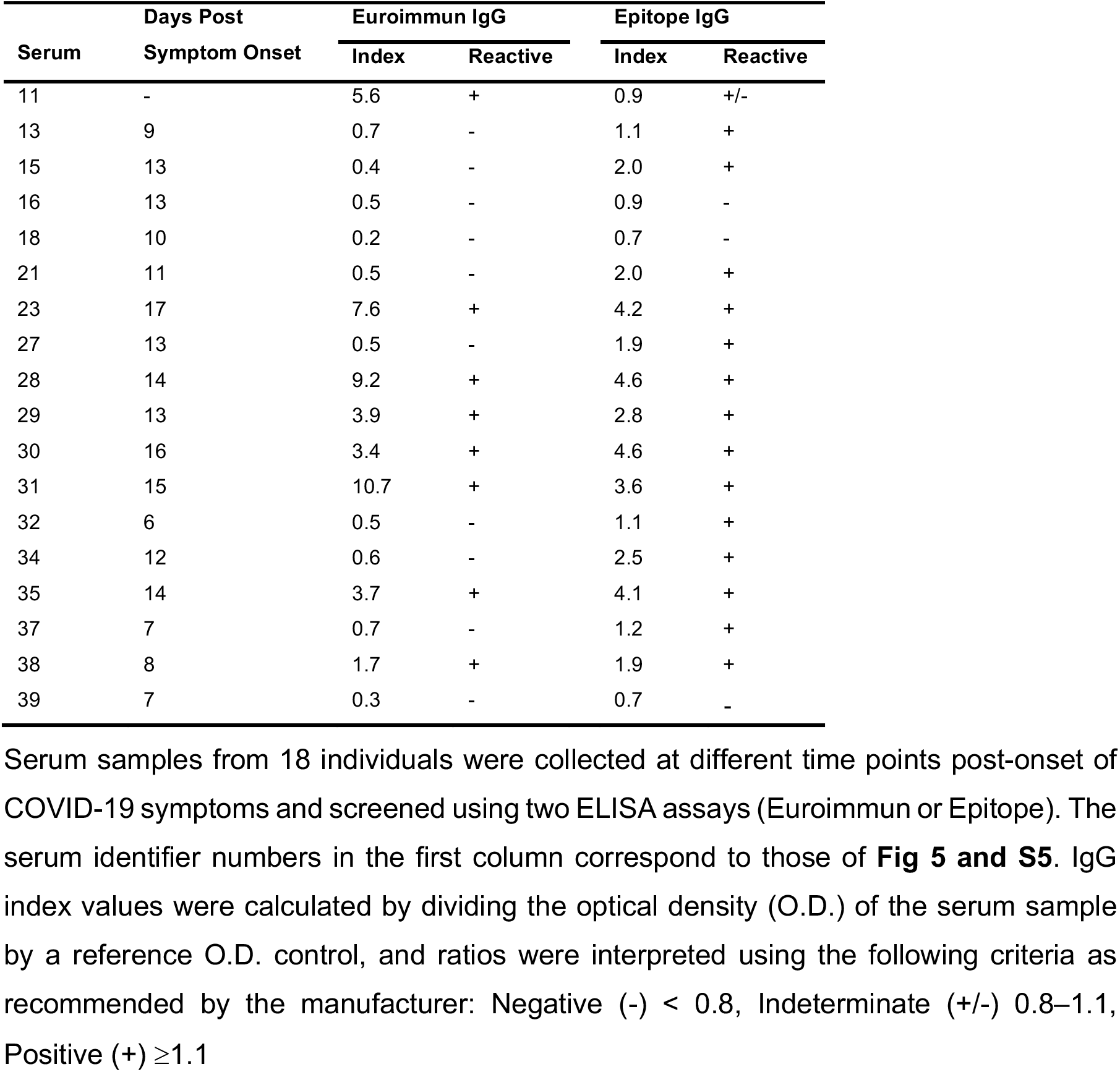
Human Serum IgG ELISA.

To extend these findings, we employed a higher throughput screening assay to test 16 additional human sera (11, 15, 16, 18, 21, 23, 27, 28, 30, 31, 32, 34, 35, 37, 38 and 39) for their ability to neutralize the VSV-SARS-CoV-2 mutants N450Y, S477N, E484A, E484D and E484K (**Table 3**). Although we observed neutralization of the various mutants at the highest concentrations of human sera tested (1:10), VSV-SARS-CoV-2 variants with substitutions at residues 484 were consistently less sensitive to neutralization by all sera (**Fig 4C and S5B**). Thus, individual escape mutants can exhibit resistance to neutralization by polyclonal human convalescent sera. This observation suggests that the repertoire of antigenic sites on RBD that bind neutralizing antibodies is limited in some humans. We again observed the increase in infectivity of substitutions at residue E484 in the presence of multiple human sera (**Fig S5B**).

### Comparison of escape mutants with S sequence variants isolated in humans

To broaden our analysis, we performed a second campaign of escape mutant selection using nine additional neutralizing mAbs generated against the RBD (**Fig 5, S6, S7 and Table 2**). This effort generated 19 additional escape mutants, bringing the total to 48. To determine whether any of the 48 escape mutants we isolated represent S protein variants circulating in humans, we compiled all publically available genome sequences of SARS-CoV-2. Using 323,183 genomes from GISAID, we calculated the substitution frequencies throughout RBD protein (**Fig 6A**) and mapped the identified residues onto RBD structure (**Fig 6B**). Of the 48 escape variants we selected, 32 are present in circulating human isolates of SARS-CoV-2 (**Fig 6A**). The most frequent S sequence variant seen in clinical isolates is D614G which is present in 69% of sequenced isolates. The fourth most frequent substitution is S477N, which is present in 4.6% of sequenced isolates and the dominant virus in Oceana. The penetrance of the remaining substitutions among clinical isolates is relatively low, with G446V, T478I, E484K, S477I and S494P ranking 79, 102, 123, 135 and 146 of the top 150 variants in S or roughly 0.05% of sequenced variants. Collectively, this analysis highlights that neutralizing mAbs against RBD can select for variants or changes at positions that already exist within the human population and establishes that some substitutions are present at high frequency.

**Figure 5.**
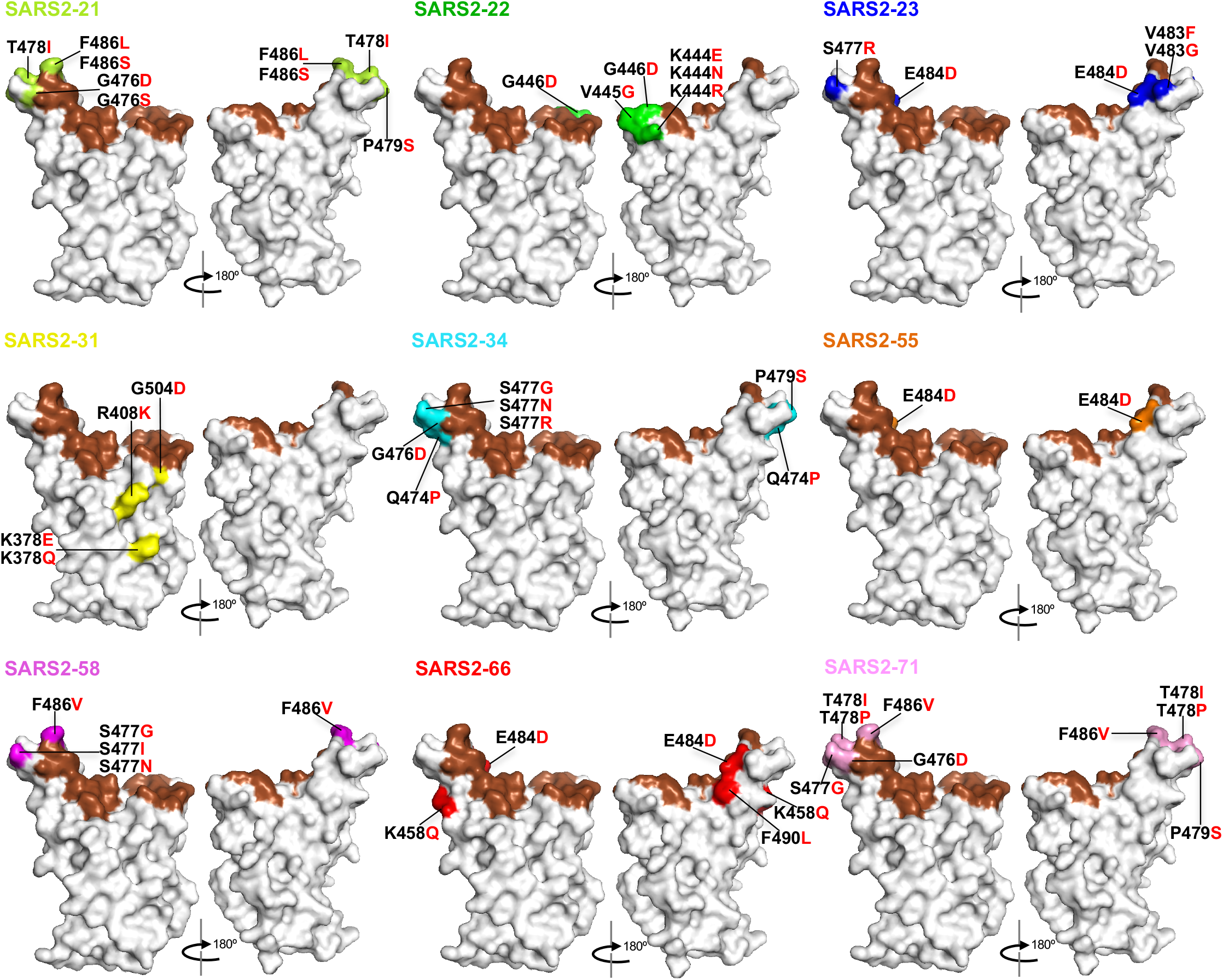
Mapping of additional VSV-SARS-CoV-2 escape mutants. The surface model of RBD (from PDB 6M0J) is depicted, and contact residues of the SARS-CoV-2 RBD-hACE2 interfaces are colored in brown. Amino acids whose substitution confers resistance to each mAb in the plaque assays are indicated for SARS2-21 (lime), SARS2-22 (green), SARS2-23 (blue), SARS2-31 (yellow), SARS2-34 (cyan), SARS2-55 (orange), SARS2-58 (magenta), SARS2-66 (red), and SARS2-71 (pink). See **Figure S6 and S7**.

**Figure 6.**
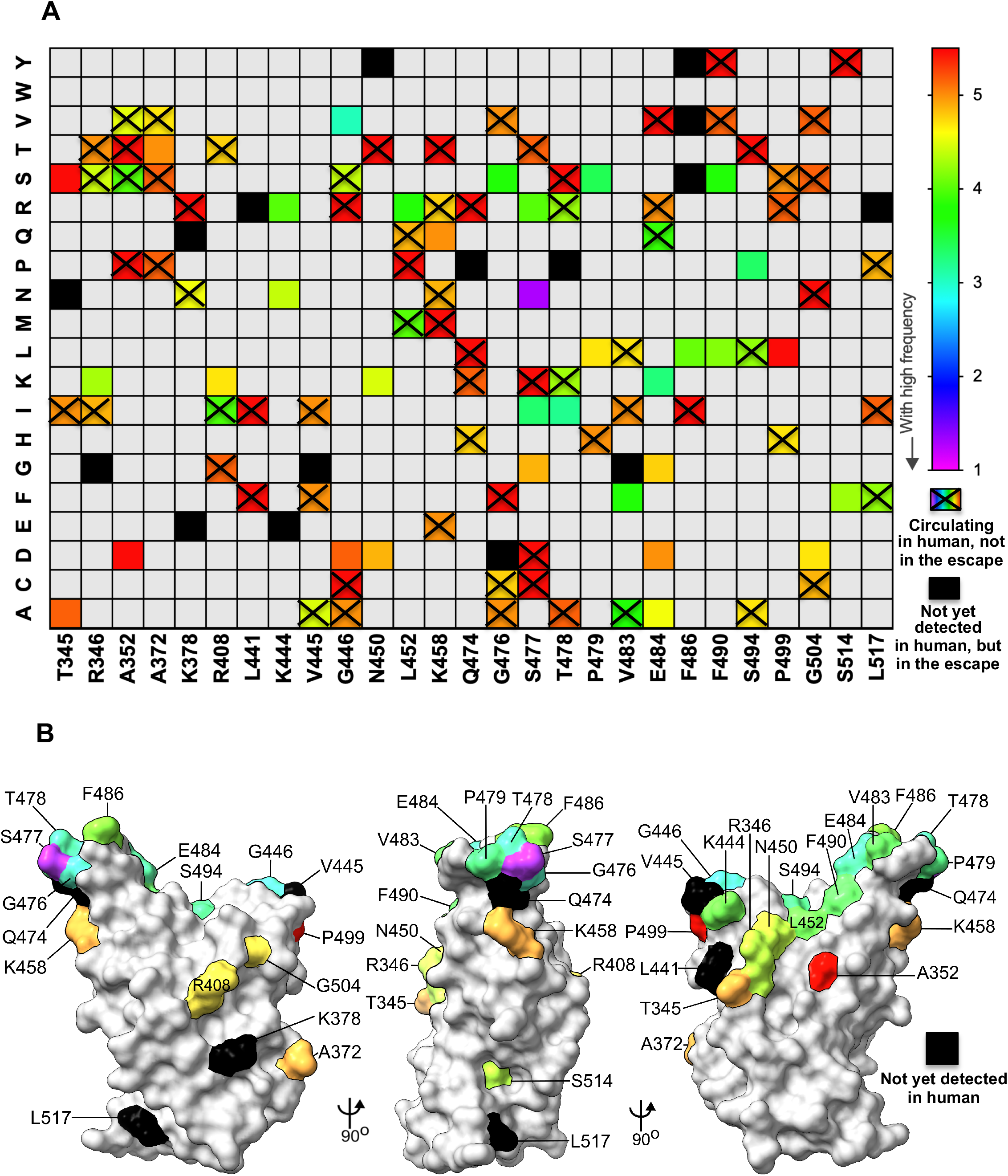
Position and frequency of RBD amino acid substitutions in SARS-CoV-2. (**A**) RBD amino acid substitutions in currently circulating SARS-CoV-2 viruses isolated from humans. For each site of escape, we counted the sequences in GISAID with an amino acid change (323,183 total sequences at the time of the analysis). Variant circulating frequency is represented as a rainbow color map from red (less circulating with low frequency) to violet (most circulating with high frequency). A black cell indicates the variant has not yet been isolated from a patient. A rainbow cell with cross indicates the variant has been isolated from a patient, but not appear in those 48 escape mutants. (**B**) Location of natural sequence variation within the RBD. The RBD is modeled as a surface representation, Variant frequency is rainbow colored as in (**A**). Black coloration indicates variation at that residue has not yet been isolated.

### Sequential selection of 2B04 and 2H04 escape mutants

To examine whether mutations resistant to antibody combinations could be isolated, we undertook a third selection campaign using a combination of 2B04 and 2H04. We were unable to isolate mutants resistant to the two antibodies when added concurrently. However, using the 2B04 resistant-viruses E484A, E484K and F486S, we selected additional mutations by growth in the presence of 2H04 (**Fig 7 and Table 2**). These selected variants were resistant to both antibodies. For the 2B04 resistant mutant E484A, we selected T345A, R346G and K444E; for mutant E484K, we isolated R346K, A372T and K444E; and for mutant F486S we selected T345S. Two of the mutants (R346K and A372T) were not seen in our prior selection campaigns with 2H04 alone, although both variants exist in human isolates (**Fig 6**). Taken together, this analysis suggests that cocktails of mAbs binding distinct epitopes on SARS-CoV-2 S protein pose an increased but not complete barrier to resistance, especially if circulating strains already encode substitutions that compromise effectiveness of one of the two mAbs.

**Figure 7.**
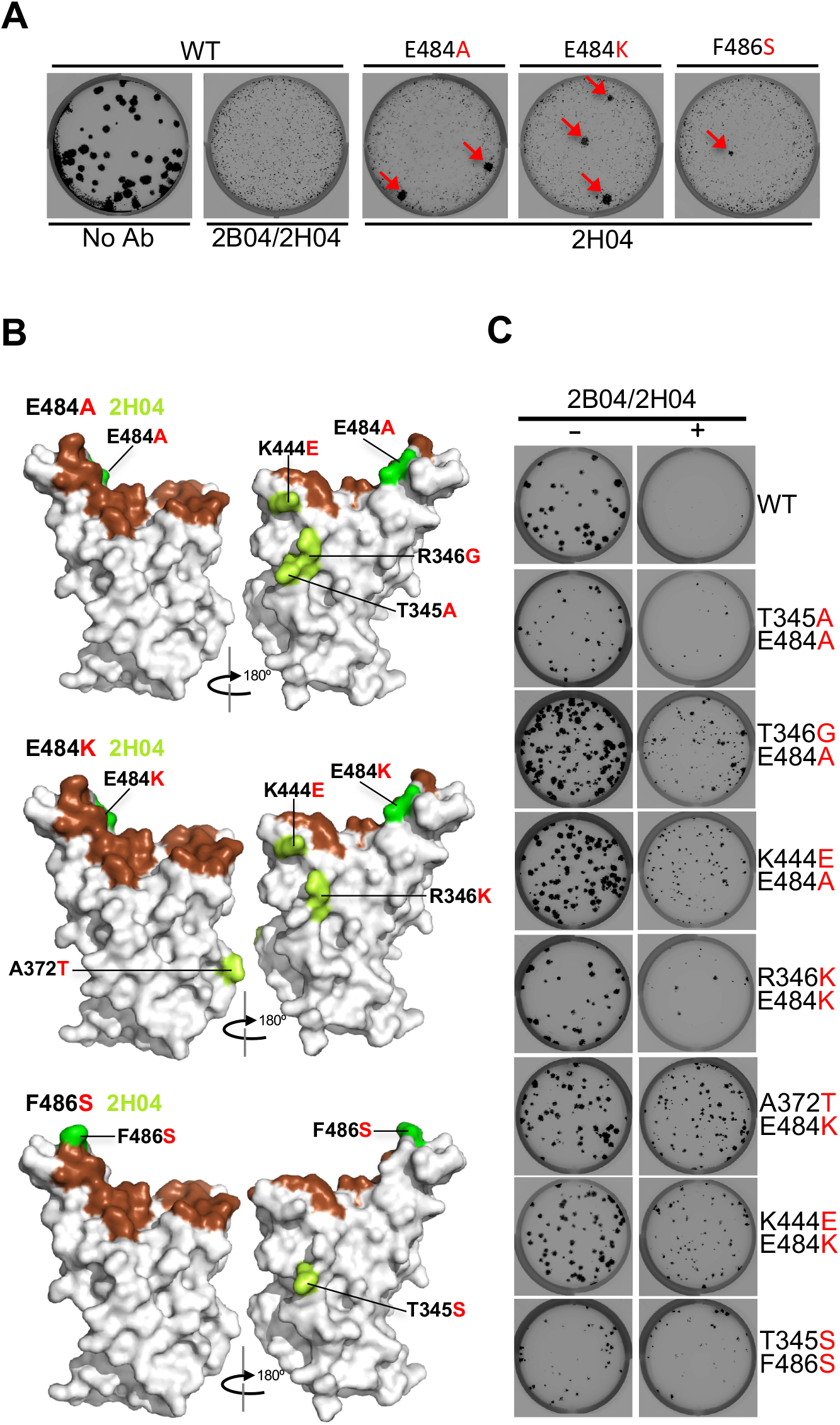
Sequential selection of 2B04 and 2H04 escape mutants. (**A**) Plaque assays were performed to isolate the VSV-SARS-CoV-2-S wild-type, E484A, E484K, and F486S escape mutant on Vero E6 TMPRSS2 cells in the present of the indicated mAb in the overlay. Representative images of three independent experiments are shown. (**B**) The surface model of RBD (from PDB 6M0J) is depicted, and contact residues of the SARS-CoV-2 RBD-hACE2 interfaces are colored in brown. 2B04 escape mutants including E484A, E484K, and F486S are indicated in green. Amino acids whose substitution confers resistance to 2H04 in the plaque assays are indicated in lemon. (**C**) Wild-type and sequentially-identified double mutants were tested for neutralizing activity using a plaque assay with the indicated mAb in the overlay. MAb concentrations added were the same as those used to select the escape mutants. Representative images of two independent experiments are shown.

## DISCUSSION

Therapeutic mAbs, convalescent plasma and vaccines are in clinical development as countermeasures against SARS-CoV-2. The efficacy of these strategies will be impacted by viral mutants that escape antibody binding and neutralization. To define the landscape of mutations in the RBD associated with resistance, we selected escape mutants to 19 neutralizing mAbs including some in clinical development. Characterization of escape mutants identified several that exhibit resistance to multiple antibodies, convalescent human sera, and soluble receptor decoys. Resistance to neutralization by serum from naturally-infected humans suggests that the neutralizing response to SARS-CoV-2 in some individuals may be dominated by antibodies that recognize relatively few epitopes. Many of the escape mutants we identified contain substitutions in residues at which variation is observed in circulating human isolates of SARS-CoV-2. If a similar limited polyclonal response occurs following S protein-based vaccination, escape variants could emerge in the human population and compromise the efficacy of such vaccines.

From 19 different mAbs that neutralize SARS-CoV-2, we isolated 50 viral mutants that escape neutralization. Selection of escape mutants was facilitated by the use of VSV-SARS-CoV-2, which we previously validated as an effective mimic of SARS-CoV-2 S protein mediated infection (Case et al., 2020). The mAbs were obtained following immunization with soluble RBD, and although some mice received a boost with stabilized S ectodomain protein, all escape substitutions map within the RBD. Multiple different mAbs led to resistance substitutions at K444, G446, N450, L452, S477, T478, P479, E484, F486 and P499, suggesting that they comprise major antigenic sites within the RBD. In earlier work, substitutions at residues K444, N450, E484 and F486 were identified using two antibodies in clinical development (Group et al., 2020), and a separate study using three different antibodies defined resistance substitutions at R346, N440, E484, F490 and Q493 (Greaney et al., 2020; Weisblum et al., 2020).

The mutations we selected also inform the mechanism by which the different antibodies function. All of the resistance mutations we identified map within or proximal to the ACE2 binding site. Likely, the majority of the antibodies we tested neutralize infection by interfering with receptor engagement. Antibodies from human survivors also interfere with receptor engagement (Wu et al., 2020b; Zost et al., 2020) suggesting a common mechanism of neutralization. Some of the resistance mutations from 2H04, SARS2-01 and SARS2-31 we identified map outside the ACE2 binding site including at the side and the base of the RBD. Direct competition with ACE2 binding is consistent with the escape mutants we selected with 2B04, whereas an indirect mechanism of action fits with the escape mutants we identified to 2H04. Our finding of an escape mutant to 2H04 located at the base of the RBD, outside the footprint of the antibody, suggests a possible allosteric mechanism of resistance. This mutation might affect the ability of the RBD to adopt the ‘up’ conformation necessary for engagement of the cellular receptors perhaps by shielding the epitope or stabilizing the RBD in the ‘down’ conformation. Further structural and functional work is required to define how different mutations promote antibody resistance and determine the mechanisms by which specific antibodies inhibit SARS-CoV-2 infection.

The relatively low genetic barrier to resistance combined with knowledge of the presence of relevant substitutions in clinical isolates suggests that effective mAb therapy likely will require a combination of at least two neutralizing antibodies (Baum et al., 2020; Du et al., 2020; Greaney et al., 2020; Group et al., 2020; Li et al., 2020; Weisblum et al., 2020). Profiling whether different residues are associated with resistance to specific antibodies could facilitate the selection of combinations based on their non-overlapping resistance mutations. Although we isolated several escape mutants that exhibit cross-resistance to multiple antibodies, other antibodies are associated with unique and non-overlapping resistance. Resistance to such combinations could still arise through sequential escape whereby a resistant variant to one antibody acquires resistance to a second. Sequential escape could be favored *in vivo* for two antibodies with different half-lives, or when a pre-existing resistant variant to one antibody already is circulating. Indeed, while we could not generate escape mutants to the antibody cocktail of 2B04 and 2H04, we readily isolated escape mutants to both mAbs through sequential selection.

Substitution S477N, the fourth most abundant S protein sequence variant in circulating human isolates of SARS-CoV-2, led to a degree of resistance to all of the mAbs we profiled, including 2B04 and 2H04. How S477N could confer such broad resistance is of interest, given its penetrance among clinical isolates (6.5%). One possible explanation may relate to changes in glycosylation at this position. Additional analysis is required to determine how broad the resistance associated with S477N is, and to probe the mechanism by which it occurs. The broad mAb resistance observed here for S477N was not accompanied by resistance to neutralization by human convalescent sera, suggesting that other epitopes – such as those centered around E484 are more dominant in humans. Among our panel of mutants, we isolated a total of 14 substitutions at sites of glycosylation, including 8 N-linked glycans site: T345N, K444N, S477N, L441R, L517R, L452R, S477R, K444R; and 6 O-linked glycans: F486S, T345S, F490S, P479S; F486Y, N450Y.

Substitutions at position E484 were associated with relative resistance to neutralization by several convalescent human sera. Four variants at this position (E484A, E484D, E484G and E484K) exhibited resistance to each of the human convalescent sera we tested. This suggests that in some humans, neutralizing antibodies may be directed toward a narrow repertoire of epitopes following natural infection. Substitution at position E484 has become increasingly common among clinical isolates. As of October 2020, just 0.03% of sequenced isolates exhibited variation at E484, which led us to suspect that variation at this position may come with an apparent fitness cost for viral replication. However, by January of 2021, the prevalence of substitutions at this position had increased to 0.09%. Substitution E484K is likely to increase in penetrance further as it linked together with N501Y and K417N changes that are present in variant 501.V2, which is believed to be more transmissible (Tegally et al., 2020). The relative resistance of the substitutions at E484 to the human sera tested highlight how variants at even a single position can affect neutralization. Given the apparent limited breadth of the human neutralizing antibody response to natural infection, it will be important to define the epitope repertoire following vaccination and develop strategies that broaden neutralizing antibody responses. In this regard, the 50 viral mutants described here, combined with additional mutants reported in related studies (Greaney et al., 2020; Group et al., 2020; Li et al., 2020; Weisblum et al., 2020), provide a compendium of functionally relevant S protein variants that could be used to profile sera from vaccine recipients in existing clinical trials.

Among the escape variants we selected, there were several that altered susceptibility to neutralization by soluble ACE2. Substitution F486S was particularly notable, as we were unable to attain 50% neutralization at the highest concentrations of soluble hACE2 tested (>20 μg/ml). The finding of an antibody escape mutant mapping to a critical residue within the ACE2 binding site raises questions regarding possible receptor usage by viruses containing S proteins with F486S. Future studies that introduce F486S into an infectious cDNA clone of SARS-CoV-2 are needed to determine the significance of this change to hACE2 interactions *in vivo*. The escape variants we selected were also examined for their sensitivity to neutralization by soluble mACE2. For the wild type S sequence and some escape mutants (*e.g*., L441R, K444N, L452R, and S477N), we observed a modest increase in infectivity at increasing concentration of soluble mACE2. Further studies using the infectious molecular clone of SARS-CoV-2 will be required to discern the significance of this observation.

We did not directly address the fitness of the mutants generated in this study, and any studies of the fitness of the VSV-SARS-CoV-2 variants would pertain to their relative replicative fitness measured in cell culture (Domingo et al., 2012). We can, however, make several inferences about fitness of specific viral mutants based on the prevalence of the corresponding mutants among circulating human isolates of SARS-CoV-2. Over 60% of the mutants we isolated in this study already circulate as natural viral variants. The escape mutants we isolated were based on single nucleotide changes starting from the sequence of the S protein of the SARS-CoV-2 Wuhan-Hu-1 strain (Wu et al., 2020a). In the context of VSV-SARS-CoV-2, the fitness of the variants in cell culture relates to their ability to resist neutralization by the indicated mAb and infect Vero cells presumably through interactions with ACE2. Our functional screens complement other systematic mutational analyses of the amino acid residues of the RBD of the SARS-CoV-2 S, such as those based on yeast display (Starr et al., 2020).

### Limitations of this study

Use of chimeric VSV that depends on SARS-CoV-2 S protein for entry into cells enabled the selection of 50 escape mutants. Although chimeric VSV serves as an effective mimic of SARS-CoV-2 S protein-mediated entry and viral neutralization, sequence analysis of circulating human isolates revealed that 34 of those escape mutants are present in the context of infectious SARS-CoV-2. The remaining 16 variants may represent S sequences with compromised fitness in the background of SARS-CoV-2 highlighting one potential limitation of our work. Additional limitations of our study are the relatively limited number of polyclonal human sera that we profiled against the panel of escape mutants. Additional human sera samples at lower dilutions may help determine the extent to which serum-based neutralization of virus is affected by individual or combinations of escape mutants.

## ACKNOWLEDGEMENTS

This study was supported by NIH contracts and grants (75N93019C00062 and R01 AI127828, R01 AI157155, and R37 AI059371) and the Defense Advanced Research Project Agency (HR001117S0019) and gifts from Washington University in Saint Louis.

## AUTHOR CONTRIBUTIONS

Z.L designed and performed the experiments. L.A.V generated and validated all hybridoma-produced mAbs. P.W.R., L.M.B., R.E.C., S.S., B. A., provided experimental assistance. M.J.L and W.J.B set up the high-throughput assay, H.Z. and D.H.F. generated and provided purified ACE2 proteins, J.M.E mapped escape mutant. E.S.T identified and provided the human immune serum. A.H.E, generated and provided cloned versions of mAbs. Z.L., M.S.D., and S.P.J.W. analyzed data. Z.L., L.A.V., M.S.D and S.P.J.W. wrote the initial draft, with the other authors providing editing comments.

## SUPPLEMENTARY FIGURE LEGENDS

**Figure S1.**
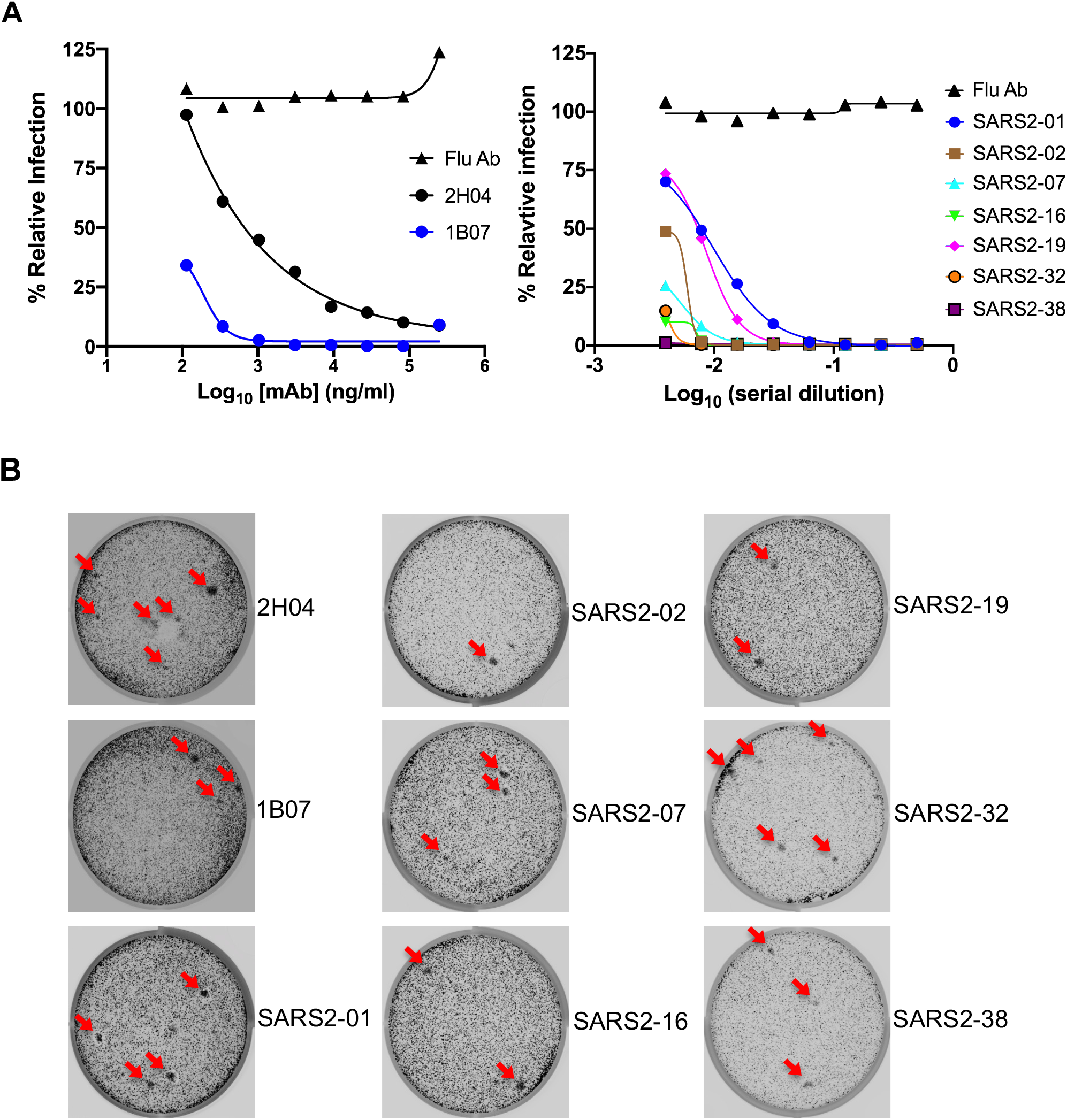
Isolation of VSV-SARS-CoV-2 escape mutants by plaque assay. Related to Fig 2. (**A**) RBD-specific antibodies were tested for neutralizing activity against VSV-SARS-CoV-2. MAbs in the left panel were purified from Expi293F cells transfected with antibody expression vector (pABVec6W) expressing heavy chain V-D-J and light Chain V-J cloned from single B cells. MAbs in the right panel were from hybridomas that bound to SARS-CoV-2-infected Vero CCL81 cells by flow cytometry. Data are representative of two independent experiments. (**B**) Plaque assays were performed to isolate the VSV-SARS-CoV-2-S escape mutant on Vero E6 TMPRSS2 cells in the present of the indicated mAb in the overlay. Representative images of two independent experiments are shown.

**Figure S2.**
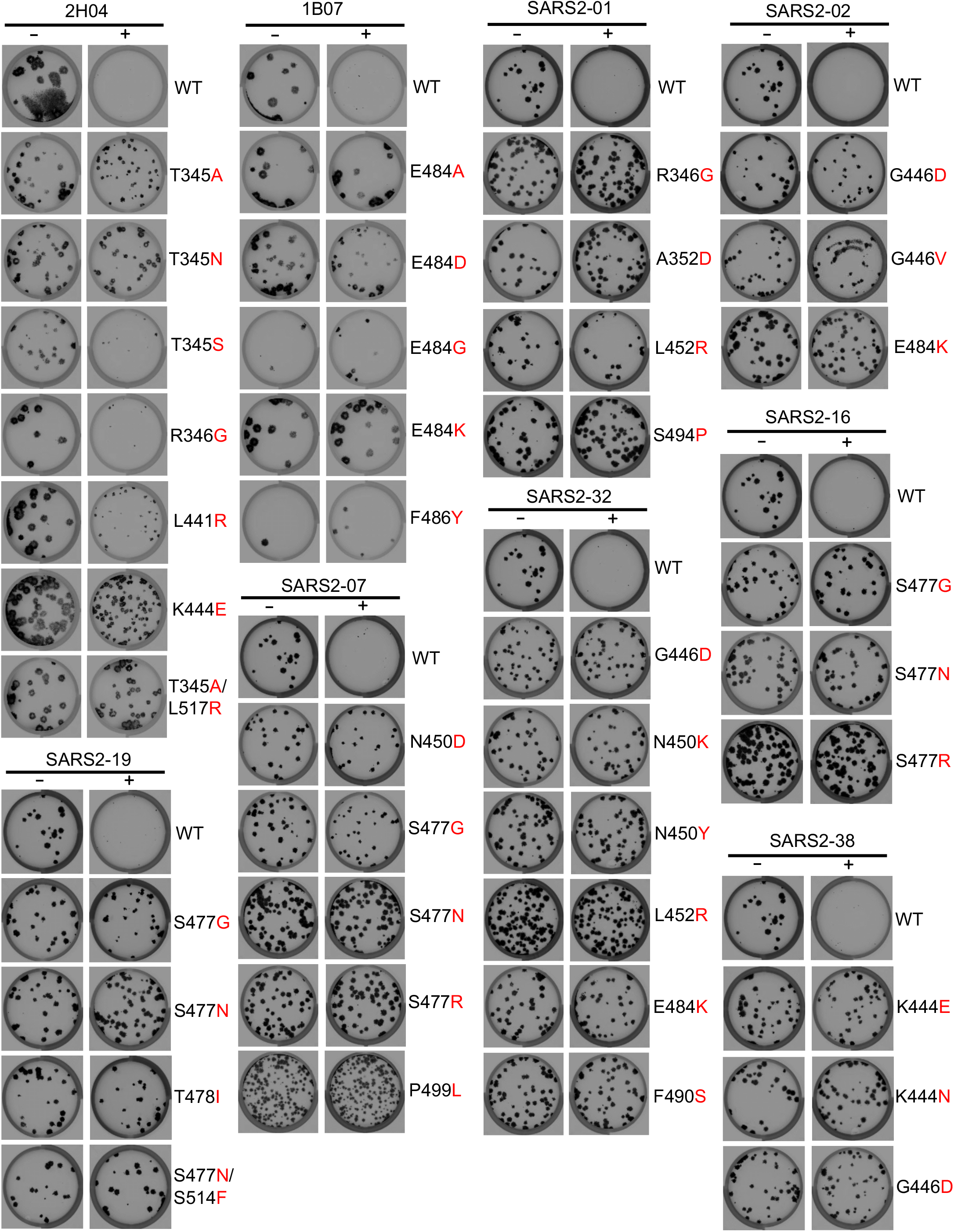
Validation of selected VSV-SARS-CoV-2 mutants. Related to Fig 2. Plaque assays were performed to validate the VSV-SARS-CoV-2 mutant on Vero cells in the presence and absence of the mAb in the overlay. MAb concentrations added in the overlay were the same as those used to select the escape mutants. Representative images of two independent experiments are shown.

**Figure S3.**
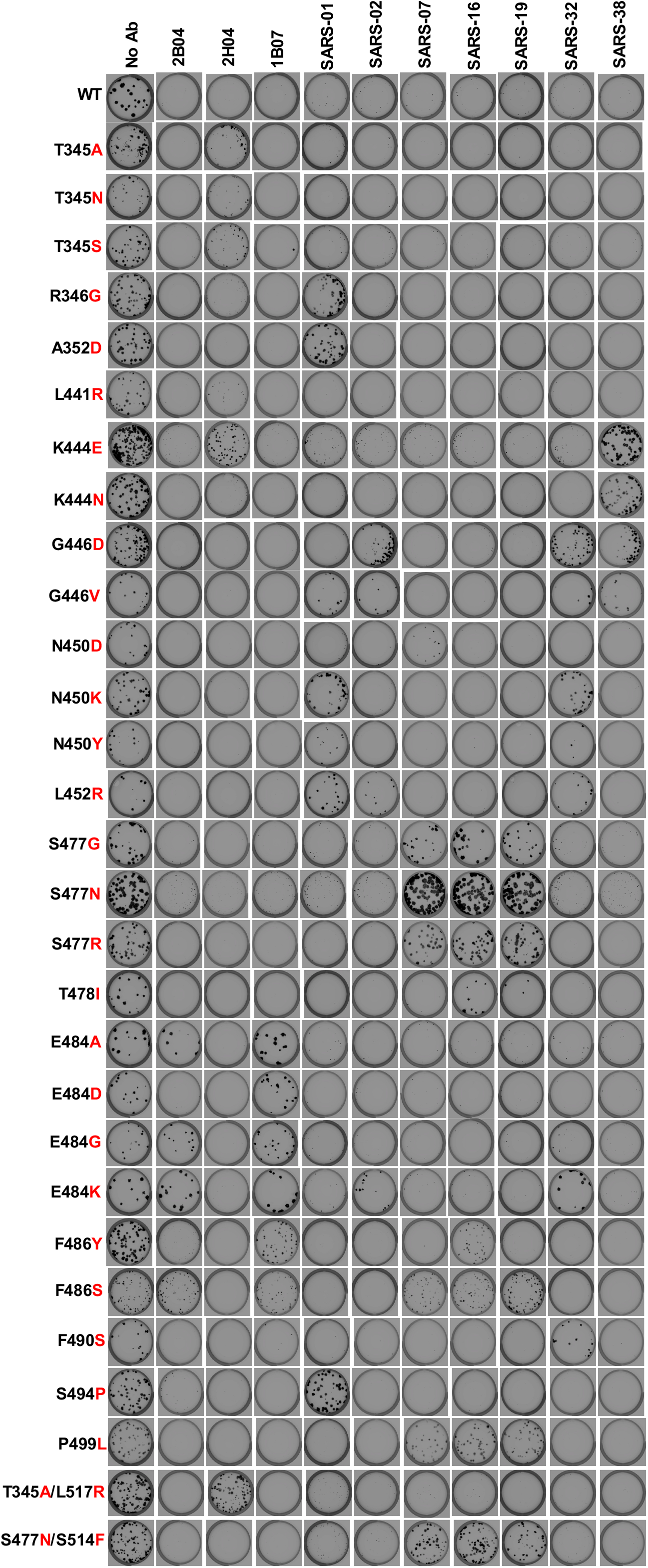
Plaque assay validation of cross-neutralization of VSV-SARS-CoV-2 mutants. Related to Fig 3A. Wild-type and identified VSV-SARS-CoV-2 mutants were tested for neutralizing activity using a plaque assay with the indicated mAb in the overlay. MAb concentrations added were the same as those used to select the escape mutants. Representative images of two independent experiments are shown.

**Figure S4.**
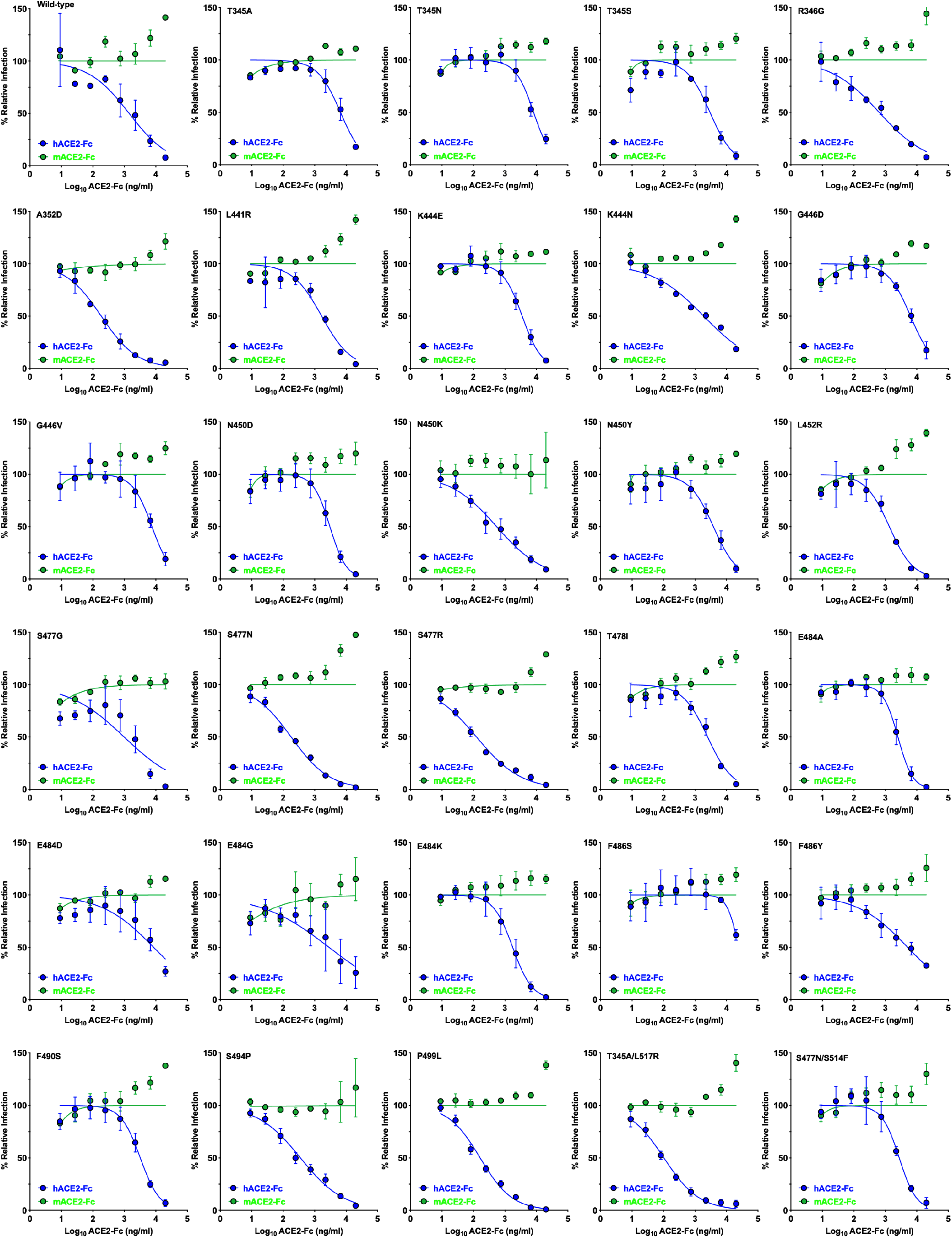
Neutralization of VSV-SARS-CoV-2 mutants by hACE2 decoy receptors. Related to Fig 3B and 3C. hACE2-Fc or mACE2-Fc were tested for neutralizing activity against wild-type and mutant VSV-SARS-CoV-2 (n=3). Error bars represent the SEM. Data are representative of three independent experiments.

**Figure S5.**
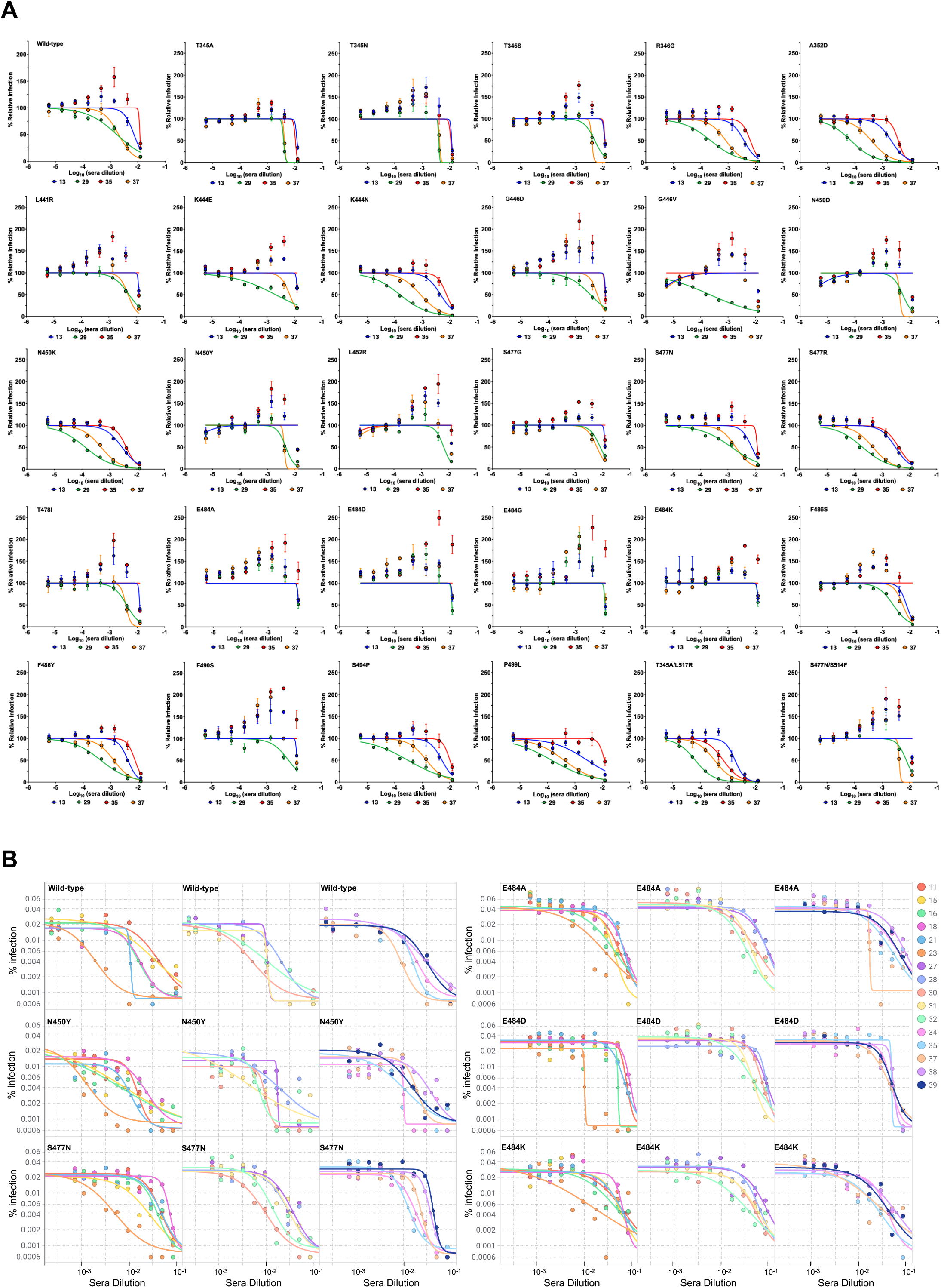
Neutralization of VSV-SARS-CoV-2 mutants by human sera. Related to Fig 4. (**A**) Four human sera were tested for neutralization of wild-type and mutant VSV-SARS-CoV-2 (n = 3). Error bars represent the SEM. Data are representative of three independent experiments. Related to **Fig 4A and 4B**. (**B**) Sixteen human sera were tested for neutralization of wild-type and 5 mutant VSV-SARS-CoV-2 (n = 1). Related to **Fig 4C**.

**Figure S6.**
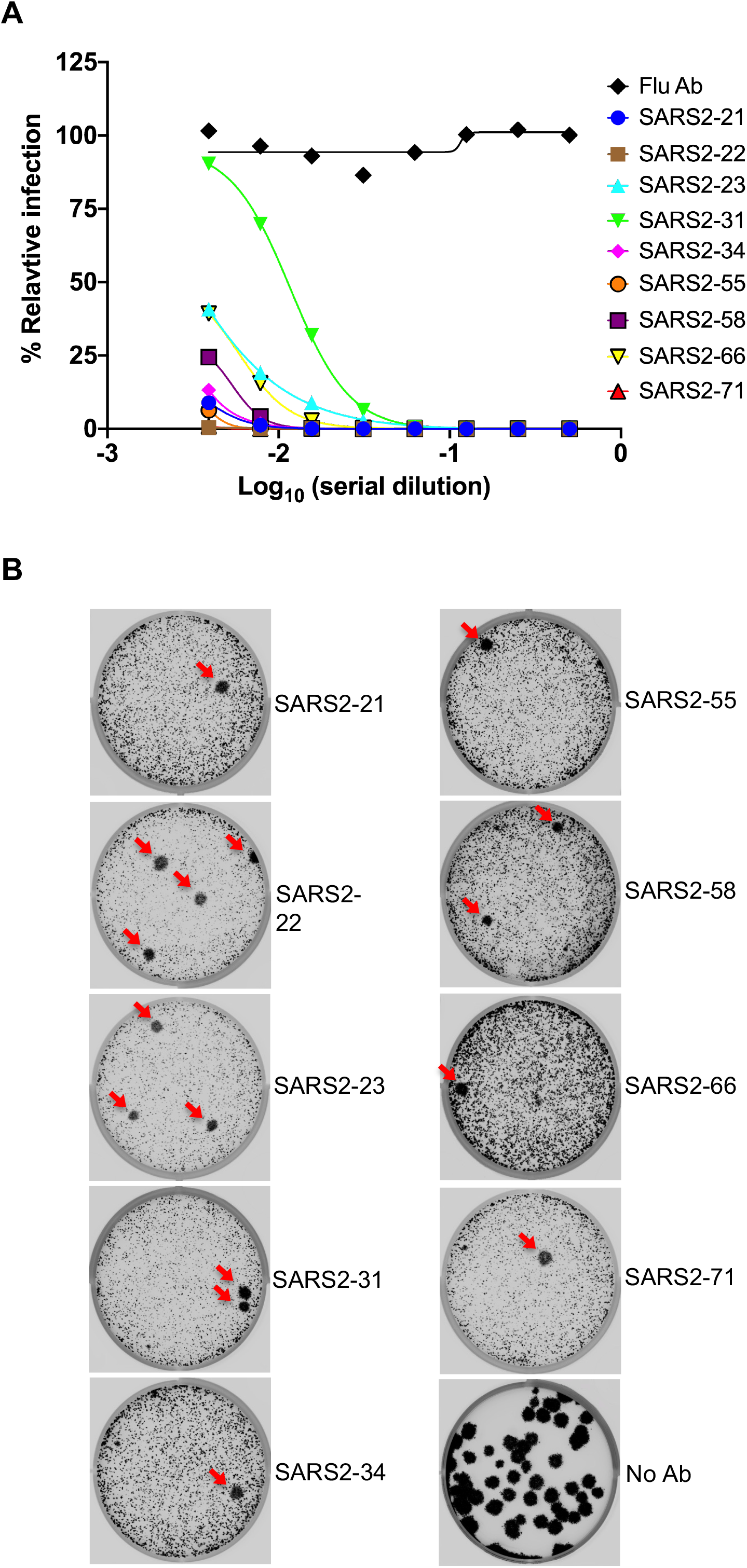
A second neutralization escape selection campaign with nine additional mAbs. Related to Fig 5. (**A**) Nine additional RBD-specific antibodies were tested for neutralization activity against VSV-SARS-CoV-2. Data are representative of two independent experiments. (**B**) Plaque assays were performed to isolate the VSV-SARS-CoV-2 escape mutant on Vero E6 TMPRSS2 cells in the presence of the indicated mAb in the overlay. Representative images of six independent experiments are shown.

**Figure S7.**
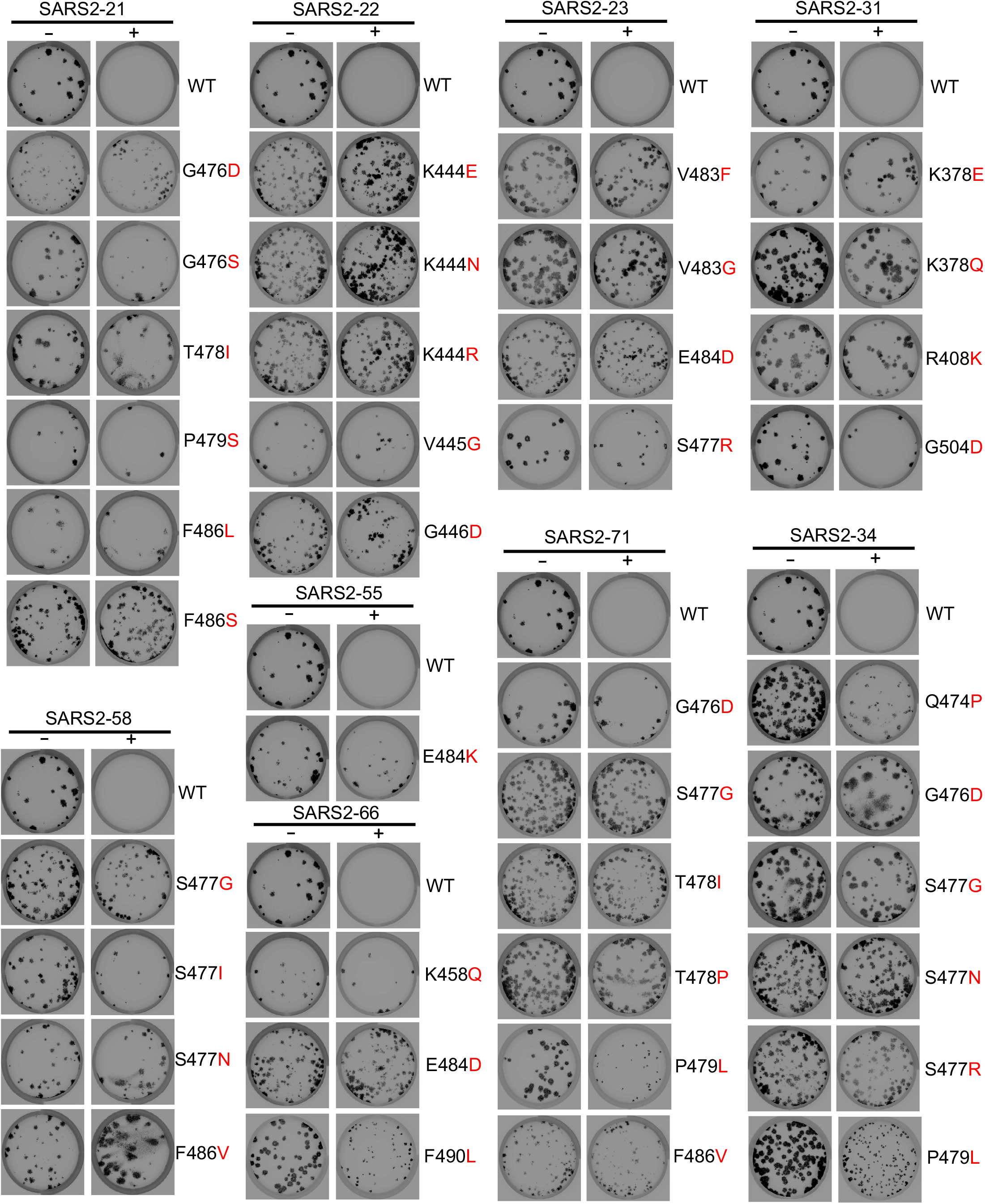
Validation of selected VSV-SARS-CoV-2 mutants. Related to Fig 5. Plaque assays were performed to validate the VSV-SARS-CoV-2 mutant on Vero cells in the presence and absence of mAb in the overlay. MAb concentration added in the overlay were the same as those used to select the escape mutants. Representative images of two independent experiments are shown.

## STAR METHODS

### RESOURCE AVAILABLITY

#### Lead Contact

Further information and requests for resources and reagents should be directed to and will be fulfilled by the Lead Contact, Sean P. J. Whelan.

#### Materials Availability

All requests for resources and reagents should be directed to and will be fulfilled by the Lead Contact author. This includes antibodies, hybridomas, viruses, and other proteins. All reagents will be made available on request after completion of a Materials Transfer Agreement.

#### Data and code availability

All data supporting the findings of this study are available within the paper and are available from the corresponding author upon request.

### EXPERIMENTAL MODEL AND SUBJECT DETAILS

#### Cells

Cells were cultured in humidified incubators at 34° or 37°C and 5% CO_2_ in the indicated media. Vero CCL81, Vero E6 and Vero E6-TMPRSS2 were maintained in DMEM (Corning or VWR) supplemented with glucose, L-glutamine, sodium pyruvate, and 10% fetal bovine serum (FBS). MA104 cells were propagated in Medium 199 (Gibco) containing 10% FBS. Vero E6-TMPRSS2 cells were generated using a lentivirus vector described as previously (Case et al., 2020).

#### VSV-SARS-CoV-2 mutants

Plaque assays were performed to isolate the VSV-SARS-CoV-2 escape mutant on Vero E6-TMPRSS2 cells with the indicated mAb in the overlay. The concentration of mAb in the overlay was determined by neutralization assays at a multiplicity of infection (MOI) of 100. Escape clones were plaque-purified on Vero-E6 TMPRSS2 cells in the presence of mAb, and plaques in agarose plugs were amplified on MA104 cells with the mAb present in the medium. Viral stocks were amplified on MA104 cells at an MOI of 0.01 in Medium 199 containing 2% FBS and 20 mM HEPES pH 7.7 (Millipore Sigma) at 34°C. Viral supernatants were harvested upon extensive cytopathic effect and clarified of cell debris by centrifugation at 1,000 x g for 5 min. Aliquots were maintained at −80°C.

#### Mouse experiments

Animal studies were carried out in accordance with the recommendations in the Guide for the Care and Use of Laboratory Animals of the National Institutes of Health. The protocols were approved by the Institutional Animal Care and Use Committee at the Washington University School of Medicine (Assurance number A3381-01). Virus inoculations were performed under anesthesia that was induced and maintained with ketamine hydrochloride and xylazine, and all efforts were made to minimize animal suffering. Female BALB/c mice (catalog 000651) were purchased from The Jackson Laboratory.

### METHOD DETAILS

#### Sequencing of the S gene

Viral RNA was extracted from VSV-SARS-CoV-2 mutant viruses using RNeasy Mini kit (Qiagen), and S was amplified using OneStep RT-PCR Kit (Qiagen). The mutations were identified by Sanger sequencing (GENEWIZ).

#### Plaque assays

Plaque assays were performed on Vero and Vero E6-TMPRSS2 cells. Briefly, cells were seeded into 6 or 12 well plates for overnight. Virus was serially diluted using DMEM and cells were infected at 37°C for 1 h. Cells were cultured with an agarose overlay in the presence of Ab or absence of Ab at 34°C for 2 days. Plates were scanned on a biomolecular imager and expression of eGFP is show at 48 hours post-infection.

#### Protein expression and purification

Soluble hACE2-Fc and mACE2-Fc were generated and purified as described as previously (Case et al., 2020).

#### Monoclonal antibodies

mAbs 2B04, 1B07 and 2H04 were described previously (Alsoussi et al., 2020). Other mAbs (SARS2-01, SARS2-02, SARS2-07, SARS2-16, SARS2-19, SARS2-21, SARS2-22, SARS2-23, SARS2-31, SARS2-32, SARS2-34, SARS2-38, SARS2-55, SARS2-58, SARS2-66 and SARS2-71) were generated as follows. BALB/c mice were immunized and boosted twice (two and four weeks later) with 5-10 μg of RBD and S protein (twice) sequentially, each adjuvanted with 50% AddaVax and given via an intramuscular route. Mice received a final, non-adjuvanted boost of 25 μg of SARS-CoV-2 S or RBD (25 μg split via intravenous and interperitoneal routes) 3 days prior to fusion of splenocytes with P3X63.Ag.6.5.3 myeloma cells. Hybridomas producing antibodies were screened by ELISA with S protein, flow cytometry using SARS-CoV-2 infected cells, and single endpoint neutralization assays.

#### Human immune sera

The human sera samples 11, 13, 15, 16, 18, 21, 23, 27, 28, 29, 30, 31, 32, 34, 35, 37, 38, 39 used in this study were previously reported (Case et al., 2020), Human donor samples were collected from PCR-confirmed COVID-19 patients. Sera samples were obtained by routine phlebotomy (Case et al., 2020). This study was approved by the Mayo Clinic Institutional Review Board.

#### Neutralization assays using a recombinant VSV-SARS-CoV-2

Briefly, serial dilutions of sera beginning with a 1:80 initial dilution were three-fold serially diluted in 96-well plate over eight dilutions. Indicated dilutions of human serum were incubated with 10^2^ PFU of VSV-SARS-CoV-2 for 1 h at 37 °C. Human serum-virus complexes then were added to Vero E6 cells in 96-well plates and incubated at 37 °C for 7.5 h. Cells were fixed at room temperature in 2% formaldehyde containing 10 μg/mL of Hoechst 33342 nuclear stain for 45 min. Fixative was replaced with PBS prior to imaging. Images were acquired using an In Cell 2000 Analyzer automated microscope (GE Healthcare) in both the DAPI and FITC channels to visualize nuclei and infected cells (×4 objective, 4 fields per well). Images were analyzed using the Multi Target Analysis Module of the In Cell Analyzer 1000 Workstation Software (GE Healthcare). GFP-positive cells were identified using the top hat segmentation method and counted within the InCell Workstation software. ACE2 neutralization assays using VSV-SARS-CoV-2 were conducted similarly. The initial dilution started at 20 μg/mL and was three-fold serially diluted in 96-well plates over eight dilutions. mAb neutralization assays using VSV-SARS-CoV-2were conducted similarly but using an MOI of 100.

#### High-throughput assay using a recombinant VSV-SARS-CoV-2

Serial dilutions of patient sera beginning with a 1:10 initial dilution were performed in 384-well plates and were incubated with 10^4^ PFU of VSV-SARS-CoV-2 for 1 h at 37°C. Vero E6 cells then were added to the human serum-virus complexes in 384-well plates at 3 × 10^3^ cells per well and incubated at 37°C for 16 h. Cells were fixed at room temperature in 4% formaldehyde and then rinsed with PBS. Cells were stained at room temperature with NucRed Live 647 (Invitrogen) for 30 min. Images were acquired using an InCell 6500 confocal imager (Cytiva) to visualize nuclei and infected cells (4X objective, 1 field per well). Images were segmented using InCarta (Cytiva). Virally-infected cells were identified by comparing to the uninfected threshold in Spotfire (Tibco). Cells were also quality-controlled (gated) based on nuclear parameters.

### QUANTIFICATION AND STATISTICAL ANALYSIS

All statistical tests were performed as described in the indicated figure legends. Non-linear regression (curve fit) was performed to calculate IC_50_ values for **Fig 3C, 4B, S4, and S5A** using Prism 8.0 (GraphPad). Non-linear regression (curve fit) was performed for **Fig 1A, S1A, and S6A** using Prism 8.0. Non-linear regression (curve fit) was performed to calculate IC_50_ values for **Fig S5B** using Spotfire (Tibco) after adding additional baseline and plateau points. Statistical significance in data **Fig 3B** was calculated by one-way ANOVA with Dunnett’s post-test using Prism 8.0. The number of independent experiments used are indicated in the relevant Figure legends.

## STAR METHODS

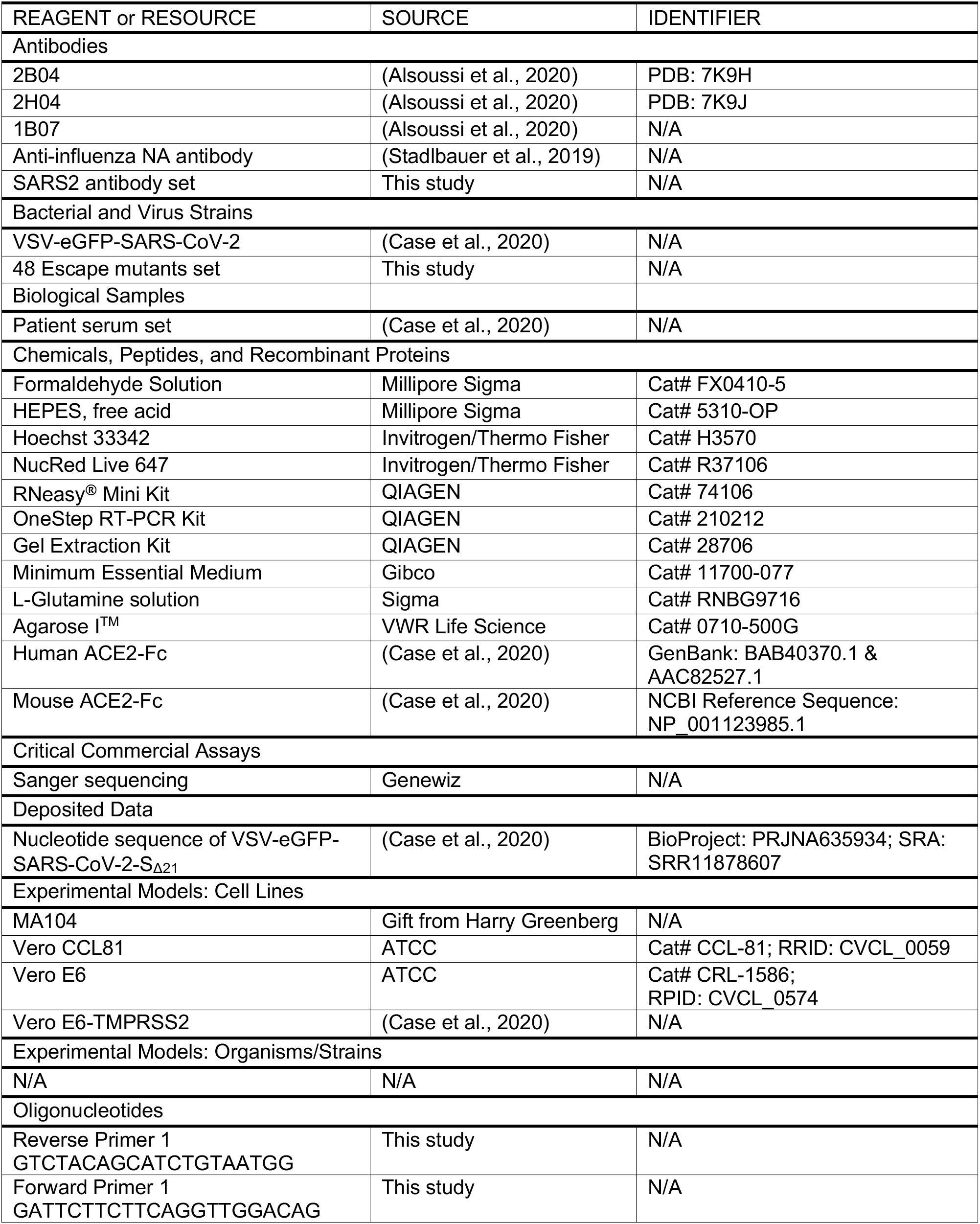
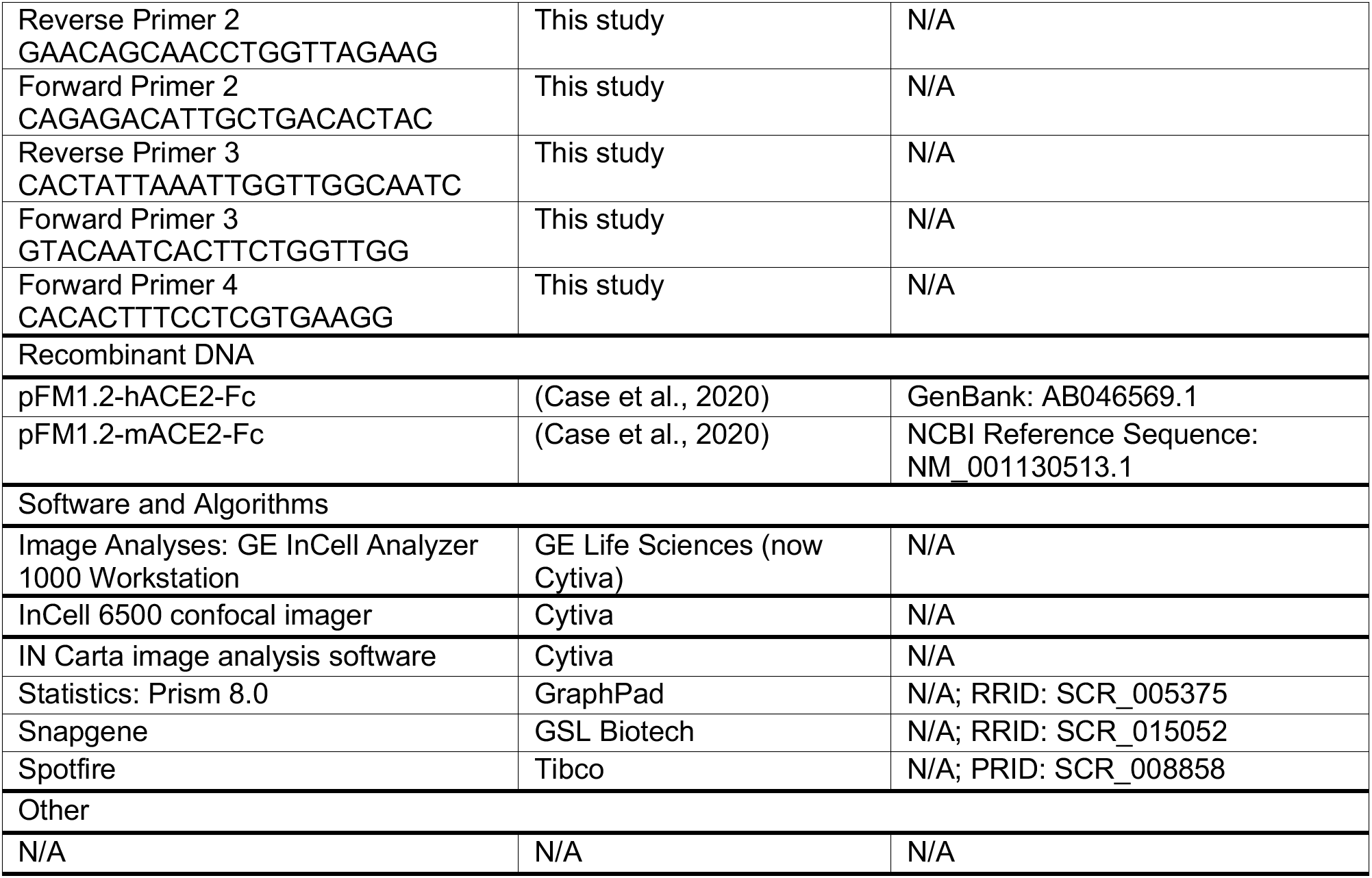
KEY RESOURCES TABLE.

